# Efficient differential expression analysis of large-scale single cell transcriptomics data using dreamlet

**DOI:** 10.1101/2023.03.17.533005

**Authors:** Gabriel E. Hoffman, Donghoon Lee, Jaroslav Bendl, N.M. Prashant, Aram Hong, Clara Casey, Marcela Alvia, Zhiping Shao, Stathis Argyriou, Karen Therrien, Sanan Venkatesh, Georgios Voloudakis, Vahram Haroutunian, John F. Fullard, Panos Roussos

## Abstract

Advances in single-cell and -nucleus transcriptomics have enabled generation of increasingly large-scale datasets from hundreds of subjects and millions of cells. These studies promise to give unprecedented insight into the cell type specific biology of human disease. Yet performing differential expression analyses across subjects remains difficult due to challenges in statistical modeling of these complex studies and scaling analyses to large datasets. Our open-source R package dreamlet (DiseaseNeurogenomics.github.io/dreamlet) uses a pseudobulk approach based on precision-weighted linear mixed models to identify genes differentially expressed with traits across subjects for each cell cluster. Designed for data from large cohorts, dreamlet is substantially faster and uses less memory than existing workflows, while supporting complex statistical models and controlling the false positive rate. We demonstrate computational and statistical performance on published datasets, and a novel dataset of 1.4M single nuclei from postmortem brains of 150 Alzheimer’s disease cases and 149 controls.

---

The human body is composed of hundreds of cell types, each with their own role in the biology of health and disease ^1^. The cell type specificity of fundamental biological mechanisms has long been appreciated ^1,2^, and genome-wide association studies have revealed strong cell type specific enrichments of risk variants that inform our understanding of disease biology ^3–5^.

Recent advances in single-cell and -nucleus transcriptomics technology have enabled profiling of over a million cells from hundreds of subjects in order to study variation in gene expression associated with disease or other traits ^6–11^. Multiplexing samples and assigning cells to subjects following sequencing, using genetic variation ^12,13^ or hashing with barcoded antibodies ^14^, has enabled a further increase in the scale of these studies. As the scale of single cell data continues to increase, and studies of cross-subject variation profile more subjects and cells, analytical workflows must keep pace.

Much of the work on differential expression analysis in single cell transcriptomics has focused on identifying expression differences between cell clusters ^15–17^. With the increasing scale of single cell datasets, recent work has examined approaches for performing differential expression analysis across subjects. Existing methods for this application were designed for small to moderate size datasets, and these model gene expression at either the single cell or the pseudobulk level for each cell type cluster ^18–21^. Modeling expression at the single cell level is the most direct approach and can be performed with a range of statistical models while allowing cell-level covariates ^18–21^. Yet statistical modeling of cell-level counts is challenging due to the low read depth and pervasive dropout effects ^18,21^, and it is essential to model the fact that multiple cells are sampled from the same subject ^19,20,22^. Even then, controlling the false positive rate remains challenging ^18^. Moreover, as studies assay more cells, the computational cost required to fit thousands of cell-level regression models becomes prohibitive ^18^.

Alternatively, pseudobulk approaches aggregate reads across cells within a cell cluster and then use methods originally developed for bulk RNA-seq ^18^. This works well for moderate sized datasets, but aggregating reads with existing methods across cells in large-scale studies can be very computationally and memory intensive. More importantly, emerging single-cell and -nucleus transcriptome datasets use complex study designs including technical or biological replicates, or use sample multiplexing that can introduce high-dimensional batch effects that existing pseudobulk methods cannot model adequately. Here, we introduce dreamlet, an open-source R package that shows superior computational and statistical performance compared to existing methods, addressing previous challenges related to statistical modeling of data from large cohorts.

## Results

### Dreamlet workflow for large-scale differential expression analysis

Dreamlet applies a pseudobulk approach and fits a regression model for each gene and cell cluster to test differential expression associated with variation in a trait across subjects. Use of precision-weighted linear mixed models enables accounting for repeated measures study designs, including technical or biological replicates, high dimensional batch effects due to sample multiplexing, and variation in sequencing depth and cell number ^23^ **(Figure 1)**. Dreamlet incorporates precision weights at two levels to account for uncertainty in the observed gene expression measurements. Dreamlet first initializes the precision weights using a Poisson count model of the observed data that considers the fact that observing more cells and reads from a sample increases the measurement precision of the underlying expression state ^18^ (**Supplementary Methods**). Incorporating these weights does not affect computational time. These initial weights are then used to estimate an empirical mean-variance trend that estimates a second round of weights without assuming a parametric model for the counting error ^24^. Dreamlet also extends the use of empirical Bayes moderated t-statistics, which borrow information across genes to increase power and control of false positive rate ^25^, to the case of precision-weighted linear mixed models (**Supplementary Methods**). Finally, the dreamlet workflow is designed for large-scale data and is substantially faster and uses less memory than existing workflows, while supporting complex statistical models and controlling the false positive rate.

**Figure 1.**
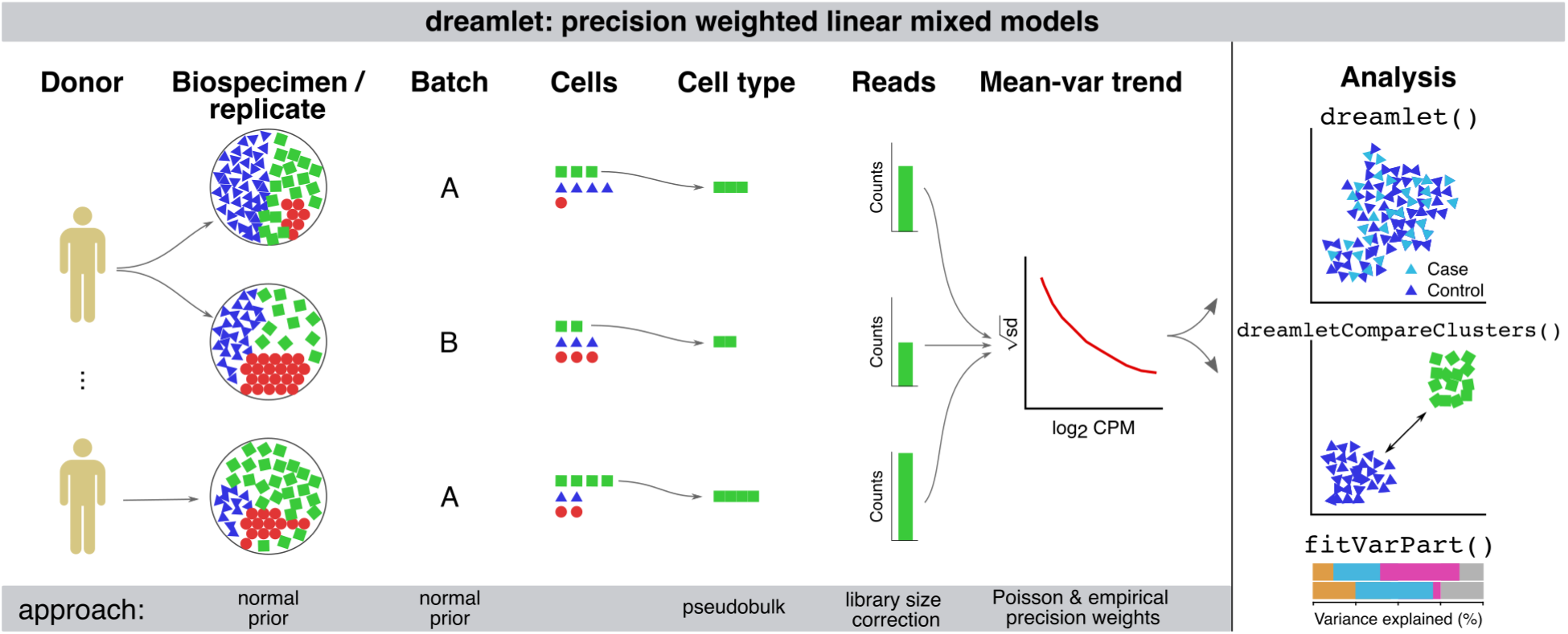
Illustration of dreamlet workflow using precision weighted linear mixed models. Expression variation across multiple biological or technical replicates, and technical batches are modeled using a random effect with a normal prior in a linear mixed model. Pseudobulk counts are computed for each cell type and standard library size correction is performed. Precision weights are initialized using an approximation of a Poisson counting model and are then used in a second weighting step to model the empirical mean-variance trend. The dreamlet package provides interfaces for differential expression analysis across donors and cell clusters, for variance partitioning analysis, and for downstream analyses and visualization.

### Computational performance on large-scale datasets

The dreamlet workflow uses an efficient implementation to compute pseudobulk counts and scales to larger datasets than competing methods. Dreamlet processes pseudobulk for 1000 donors across 12 cell types for 2.2 million cells in 10 minutes using only 20 Gb of memory (**Figure 2A,B**). Performance of pegasus ^26^ is slightly faster but its memory footprint is substantially larger and increases rapidly with sample size. Other methods are 10-100x slower and are limited by memory constraints even for moderate sample sizes.

**Figure 2:**
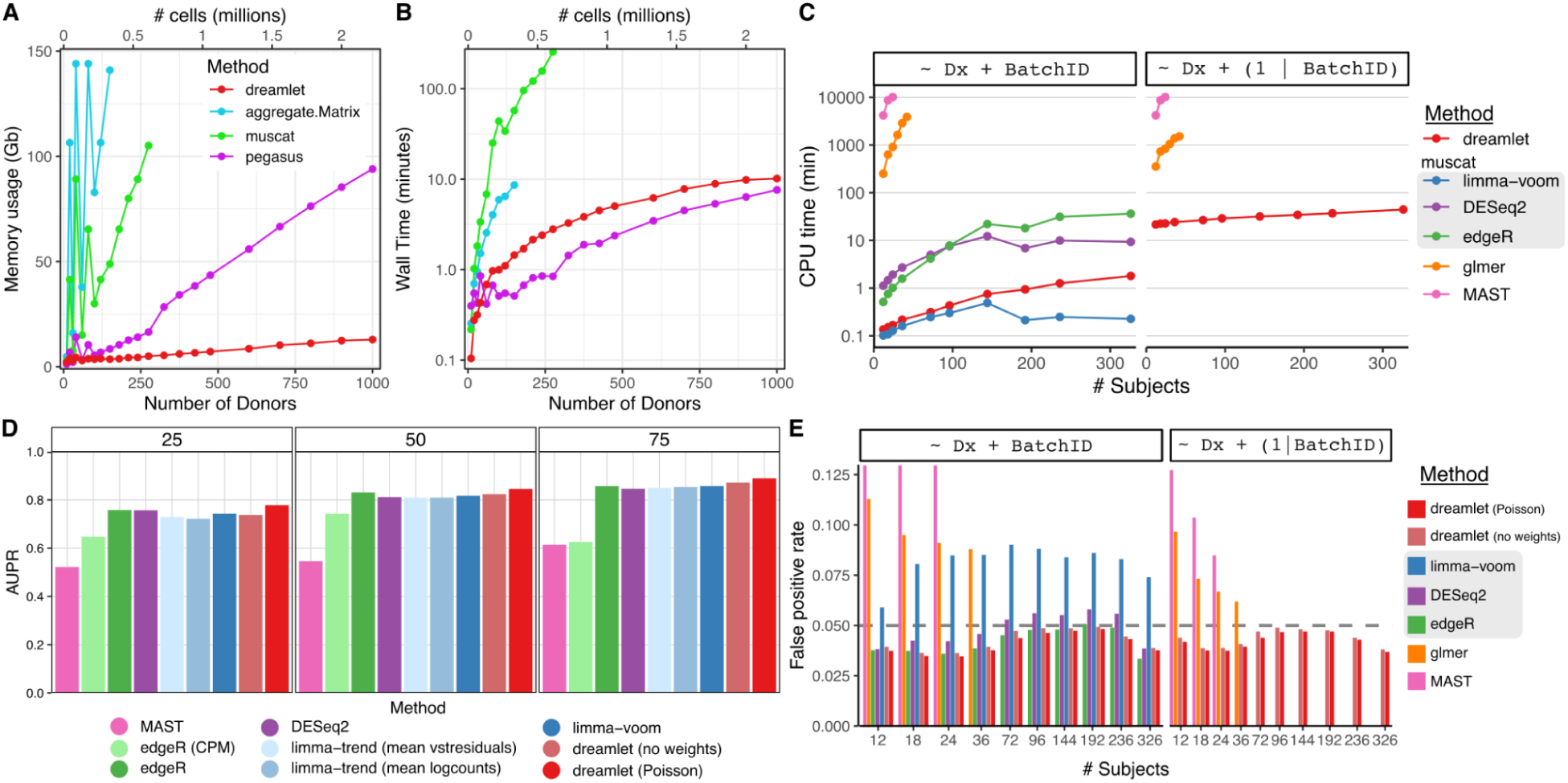
Computational and statistical performance of dreamlet workflow. **A,B)** Peak memory usage (**A**) and CPU time (**B**) averaged across 10 runs using 1 CPU core on a machine with 144 Gb memory. (**C**) CPU times for differential expression analysis for an increasing number of subjects for 6 differential expression methods. Methods in the gray box use a pseudobulk approach run using muscat software, while glmer and MAST model the data at the single cell level using generalized linear (mixed) models. (**D**) Performance of 9 differential expression methods with sample size increasing from 25 to 75 subjects measured by area under the precision recall curve (AUPR). (**E**) False positive rates for an increasing number of subjects using real single nucleus RNA-seq from postmortem human brains and permuting the disease status. Dashed horizontal gray lines indicate the target false positive rate of 5%. Results are shown for testing the effect of diagnosis, while modeling batch as a fixed (left) or random effect (right). Methods in the gray box use a pseudobulk approach run using muscat software, while glmer and MAST model the data at the single cell level.

Dreamlet fits precision-weighted linear mixed models for each gene parallelized across multiple CPU cores and enables analysis of 12 cell types across 326 subjects in 45 CPU minutes corresponding to a wall time of 15 minutes in this case (**Figure 2C**). This is over an order of magnitude faster than using generalized linear mixed models (GLMMs) at the single cell level, and competitive with other pseudobulk approaches using a negative binomial model that are not able to model random effects (i.e. edgeR ^27^, DESeq2 ^28^).

### Statistical performance on large-scale datasets

Using a simulation pipeline and benchmarks developed by an independent group ^18^, dreamlet using only a fixed effects model shows statistical performance matching or exceeding the best performing current methods while giving the most accurate estimates of the simulated effect sizes (**Figure 2D, Supplementary** Figures 1-2). Using the initial weights from a Poisson model boosts performance of dreamlet compared to the unweighted version. We note that these simulations used a simple study design where all 9 differential expression methods were applicable, while dreamlet is the only scalable method that can model complex study designs using random effects. Yet simulating single cell transcriptomics data that accurately recapitulates the complexity of real datasets is notoriously challenging ^18,29^. Instead, we used real data from human postmortem brains and permuted the disease status to estimate the empirical false positive rate across differential expression methods within each of 12 cell types. Of the methods that control the false positive rate across a range of sample sizes, dreamlet is the only one that can model random effects (**Figure 2E**). Results are similar when examined within each of the 12 cell types. Notably, the limma/voom method applied by muscat ^18^ shows inflated false positive rates in most conditions. Generalized linear mixed models used by the glmer ^30^ and MAST ^21^ methods can model random effects, but have an inflated false positive rate for small and moderate sample sizes. This is consistent with the fact that the null distributions of the coefficient estimates for these methods are normally distributed only in the asymptotic limit of large sample size. Additional methods tested by others do not scale to these large datasets ^18–20^.

Multiplexing combines multiple samples in parts of the experimental workflow and reduces the cost of generating large-scale datasets. This process creates many small batches, each of 6 to 12 samples, that can share technical artifacts. Statistically, these batches are termed ‘high dimensional’ when their number is large compared to the sample size. In order to control the false positive rate, a statistical model should account for these high dimensional batch effects. Yet widely used fixed effects models can suffer from a substantial decrease in power, and can even perform worse when including the batch effect compared to omitting it from the model ^31^. We show that using a linear mixed model to account for batches with a random effect controls the false positive rate while retaining power (**Supplemental Figure 3**).

Next we apply dreamlet to single cell RNA-seq from COVID-19 patients, and bone metastasis from prostate cancer. We then describe a novel single nucleus RNA-seq data from an Alzheimer’s disease cohort, and perform analysis using dreamlet to examine cell type specific biology of the disease.

### Dreamlet uncovers cell type specific response associated with COVID-19 severity

The COMBAT Consortium ^32^ collected blood from patients hospitalized with COVID-19 or sepsis, along with blood from non-hospitalized COVID-19 patients and healthy controls. We performed dreamlet analysis on single cell transcriptome data of 674K cells from 110 donors to identify transcriptional response to SARS-CoV2 infection associated with COVID-19 severity. The immune response associated with infection status is substantial and varies across cell types (**Supplementary** Figure 4), so we focus on monocyte subtypes and related cell types due to their key role in inflammatory response (**Figure 3A**). Analysis of pseudobulk counts for classical monocytes shows a strong mean-variance trend in the log_2_ counts which dreamlet models using two-step precision weights (**Figure 3B**). Variance partitioning analysis for each gene in classical monocytes estimates the fraction of expression variance attributable to variation across 6 disease states (i.e. 4 COVID-19 severity levels, sepsis, and healthy control), age and sex (**Figure 3C**). Expression variation across disease states was the strongest source of variation with a median of 11.4% and 3,250 of the 12,472 genes with sufficient expression explaining more than 25% of expression variance. Genes show a number of variance partitioning profiles with *CLU* showing variation across disease states, PRDX2 showing variation across age, and *XIST* varying across sex since it is on the X chromosome (**Figure 3D**). Comparing gene expression in classical monocytes between patients with mild COVID-19 and healthy controls identified 1,824 differentially expressed genes (**Figure 3E**). Key response genes showed different expression patterns, with *CLU* expression increasing with COVID-19 severity and sepsis compared to healthy controls while *NFKBIA* upregulated in COVID-19 but not sepsis patients (**Figure 3F**). Gene set analysis using the full spectrum of test statistics ^33^ identified key immune response pathways activated in different cell types based on disease state (**Figure 3G**). TNFα signaling via NF-κB showed activation in all COVID-19 disease states but not in patients with sepsis, while cholesterol homeostasis, interferon-γ and oxidative phosphorylation were activated in hospitalized COVID-19 patients but not those that were not admitted. Within the TNFα signaling pathway genes showed different patterns based on disease state, with *NFKBIA* upregulation specific to COVID-19 patients, *TNF* activated only in mild and non-hospitalized COVID-19 patients, and *CD83* unregulation specific to non-hospitalized COVID-19 patients (**Figure 3H**).

**Figure 3:**
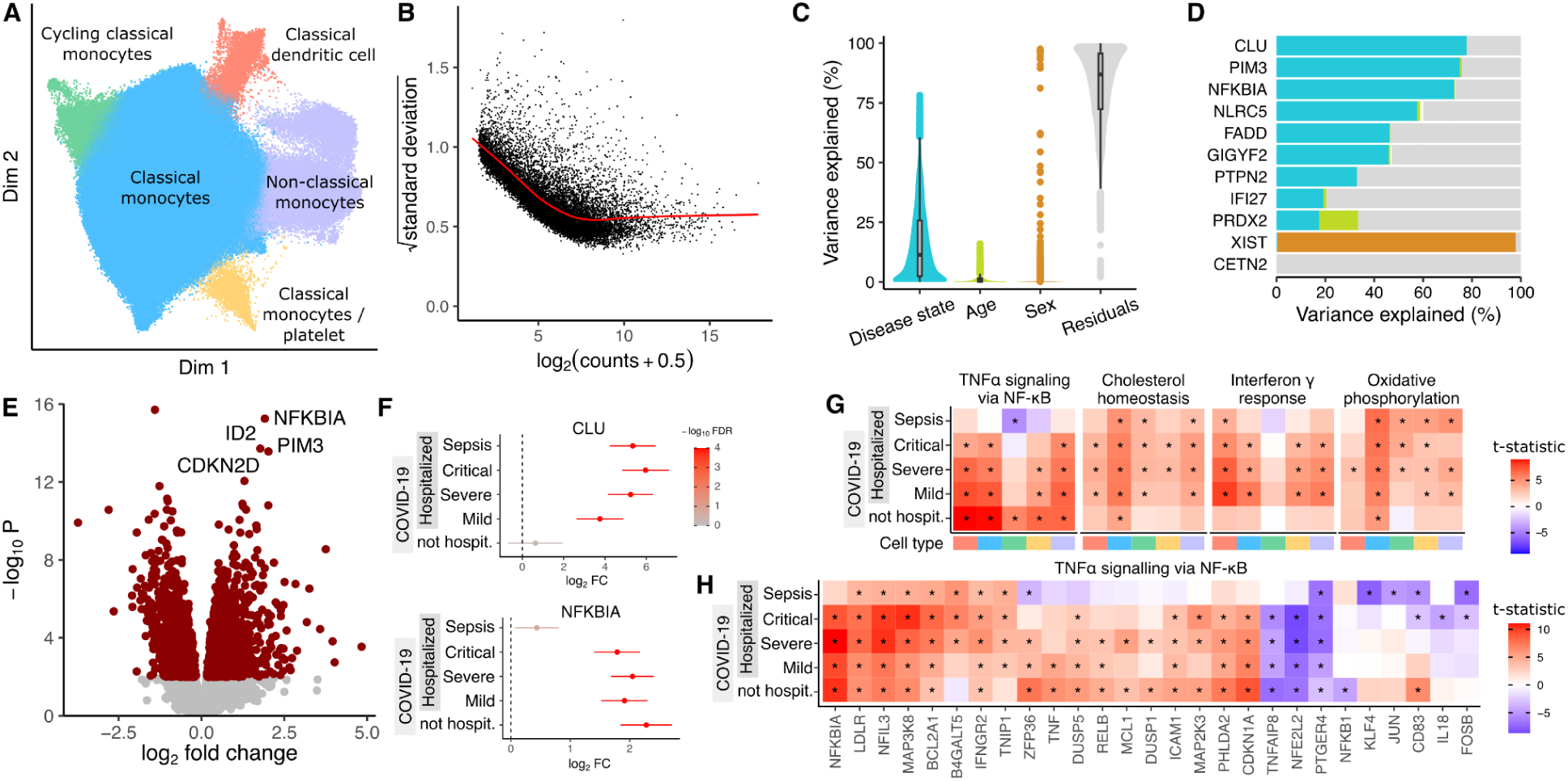
Dreamlet analysis of expression response associated with COVID-19 severity in monocyte populations. **A)** Dimensionality reduction showing monocyte subtypes and related cell types. **B)** Plot of mean-variance trend for expression counts in classical monocytes**. C)** Violin plot summarizing variance partitioning analysis of classical monocytes separating the fraction of expression variation for each gene into 4 components. **D)** Representative genes with high variance fractions explained by each of the 4 components. **E)** Volcano plot of differential analysis between mild COVID-19 and healthy controls. Red points indicate genes with FDR < 5%. **F)** Forest plot showing log_2_ fold change of expression of *CLU* and *NFKBIA* in disease states compared to healthy controls. Color indicates false discovery rate after study-wide correction for multiple testing. Error bars indicate 95% confidence interval. **G)** Heatmap of test statistics from gene set enrichment analysis shown for 5 disease states compared to healthy controls across 5 cell subtypes. Cell type color is indicated in panel (**A**). Study-wide FDR < 5% is indicated by ‘*’. **H)** Heatmap of test statistics for a subset of genes in the gene set ‘TNFα signaling via NF-κB’ from MSigDB ^34^.

### Dreamlet identifies robust transcriptional changes in bone metastases from prostate cancer

Prostate cancer can metastasize to the bone and result in very poor patient prognosis. Kfoury et al. ^7^ performed single cell RNA-seq on solid tumor, involved bone marrow, and distal bone marrow from 9 prostate cancer patients with bone metastases, as well as benign bone marrow from 7 patients without cancer. We applied the dreamlet workflow to identify genes that were differentially expressed based on disease status by modeling the multiple measurements from each cancer patient using a random effect. The number of expressed and differentially expressed genes varied widely across cell types and group comparisons (**Supplementary** Figure 5). Analysis of pseudobulk counts for a monocyte cluster (**Figure 4A**) shows the typical mean-variance trend that dreamlet models (**Figure 4B**). Variance partitioning analysis in this cell type estimates the fraction of expression variance attributable to variation across patients, disease status of each sample (i.e., tumor, involved bone marrow, etc), and disease status of the patient (i.e. cancer vs non-cancer). Expression variation across patients is the strongest, while disease status explains greater than 10% of expression variation for 525 of 1899 (27.6%) genes with sufficient expression to be included in the analysis (**Figure 4C**). This is consistent with cancer being a much stronger driver of gene expression changes than TB, above. Genes show a range of variance partitioning profiles (**Figure 4D**). For example, while *CCNL1* shows high variance across disease status, *VIM* has high variation explained by patient status (**Figure 4E**). Dreamlet analysis identified 157 genes in this monocyte cluster as differentially expressed between tumor and involved bone marrow at a study-wide FDR of 5% (**Figure 4F**). Gene set analysis using the full spectrum of test statistics identified upregulation of MHC class II proteins and protein folding chaperones in the tumor samples across a range of cell types, including multiple monocyte clusters (**Figure 4G**). Examining protein folding chaperones shows upregulation in tumor samples for most genes with sufficient expression (**Figure 4H**).

**Figure 4:**
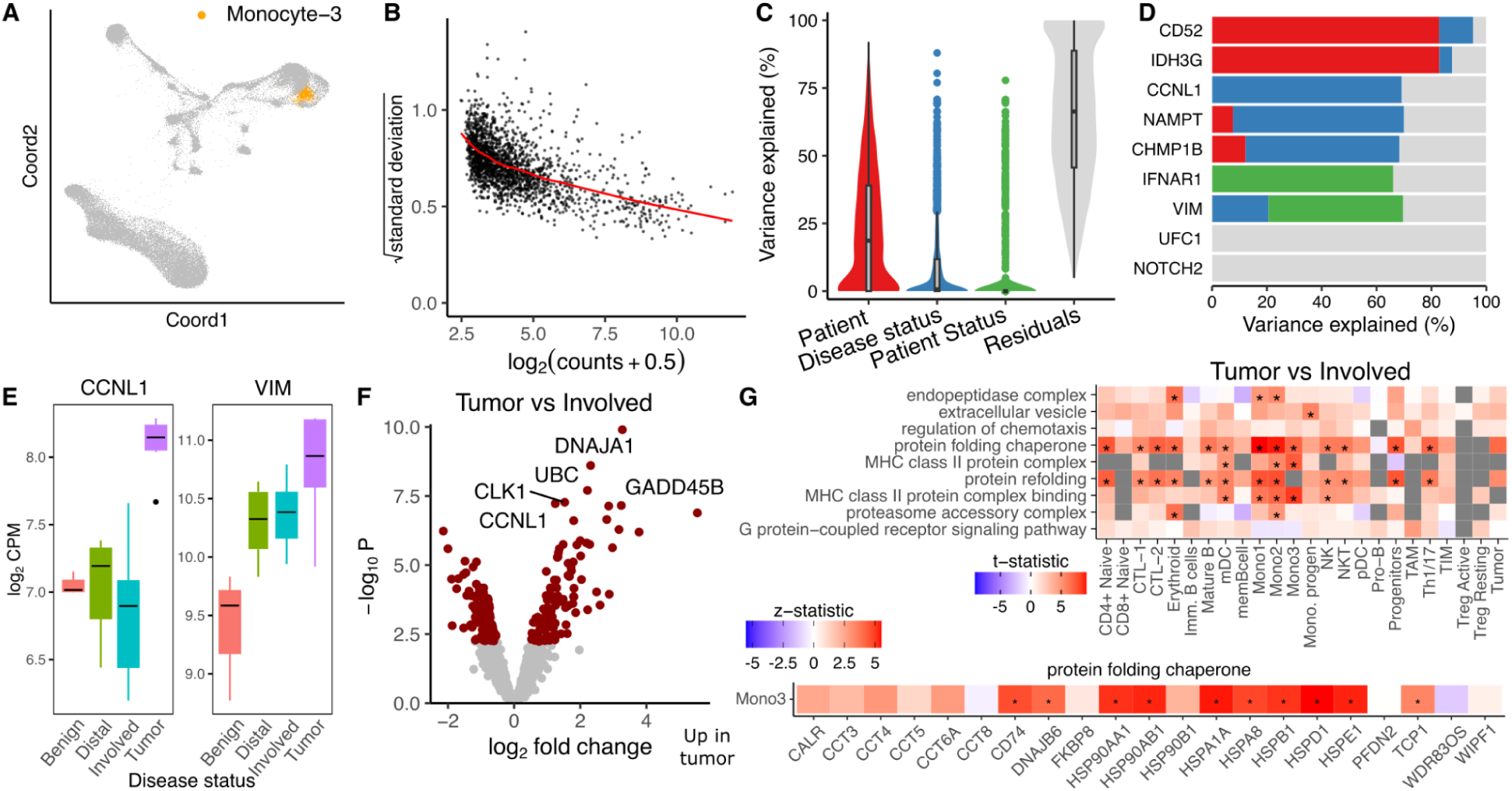
Dreamlet analysis of expression differences in prostate cancer bone metastases. **A)** Dimensionality reduction highlighting monocyte population. **B)** Plot of mean-variance trend for expression counts in this monocyte population. **C)** Violin plot summarizing variance partitioning analysis separating the fraction of expression variation for each gene into 4 components. **D)** Representative genes with high variance fractions explained by each of the 4 components. **E)** Representative genes with high expression variation across disease state (*CCNL1*) and subject cancer status (*VIM*). **F**) Volcano plot of differential analysis between tumor and involved bone marrow within each subject. **G)** Gene set analysis using the full spectrum of test statistics shows cell type specific signatures of tumor vs involved bone marrow. Study-wide FDR < 5% is indicated by ‘*’.

### Modeling multiple sources of expression variation in a large Alzheimer’s Disease cohort

We generated single nucleus RNA-seq (snRNA-seq) on tissue samples from the dorsolateral prefrontal cortex (DLPFC) of postmortem brains from donors in the Mount Sinai NIH NeuroBioBank (**Figure 5A**). Samples were multiplexed by pooling 6 donors using the nuclei hashing method ^35^. Each pool was processed in duplicate using the 10x Genomics single cell gene expression platform, to produce technical replicates. Following raw data processing and quality control, the dataset comprised 586 samples from 299 donors over 60 years of age and 1.4M nuclei. Of these donors, 150 had Alzheimer’s disease (AD) and 149 were age-matched neurotypical controls (**Supplementary** Figure 6). Cell cluster annotation identified 22 cell types including 8 subtypes of excitatory neurons and 6 subtypes of inhibitory neurons (**Figure 5B**). Analysis of pseudobulk counts for microglia (Micro_PVM) shows the strong mean-variance trend (**Figure 5C**) that dreamlet models with precision weights. Variance partitioning analysis for each gene in this cluster estimates the fraction of expression variance attributable to variation across technical replicates from the same subject, as well as variation across age, AD status, sample pool and sex (**Figure 5D**). This identifies genes with different variance profiles where, for example, 20.2% of the variation in *PTPRG* is explained by Alzheimer’s status while 97.3% of variance in NXPE1 is explained by sample pool (**Figure 5E**).

**Figure 5:**
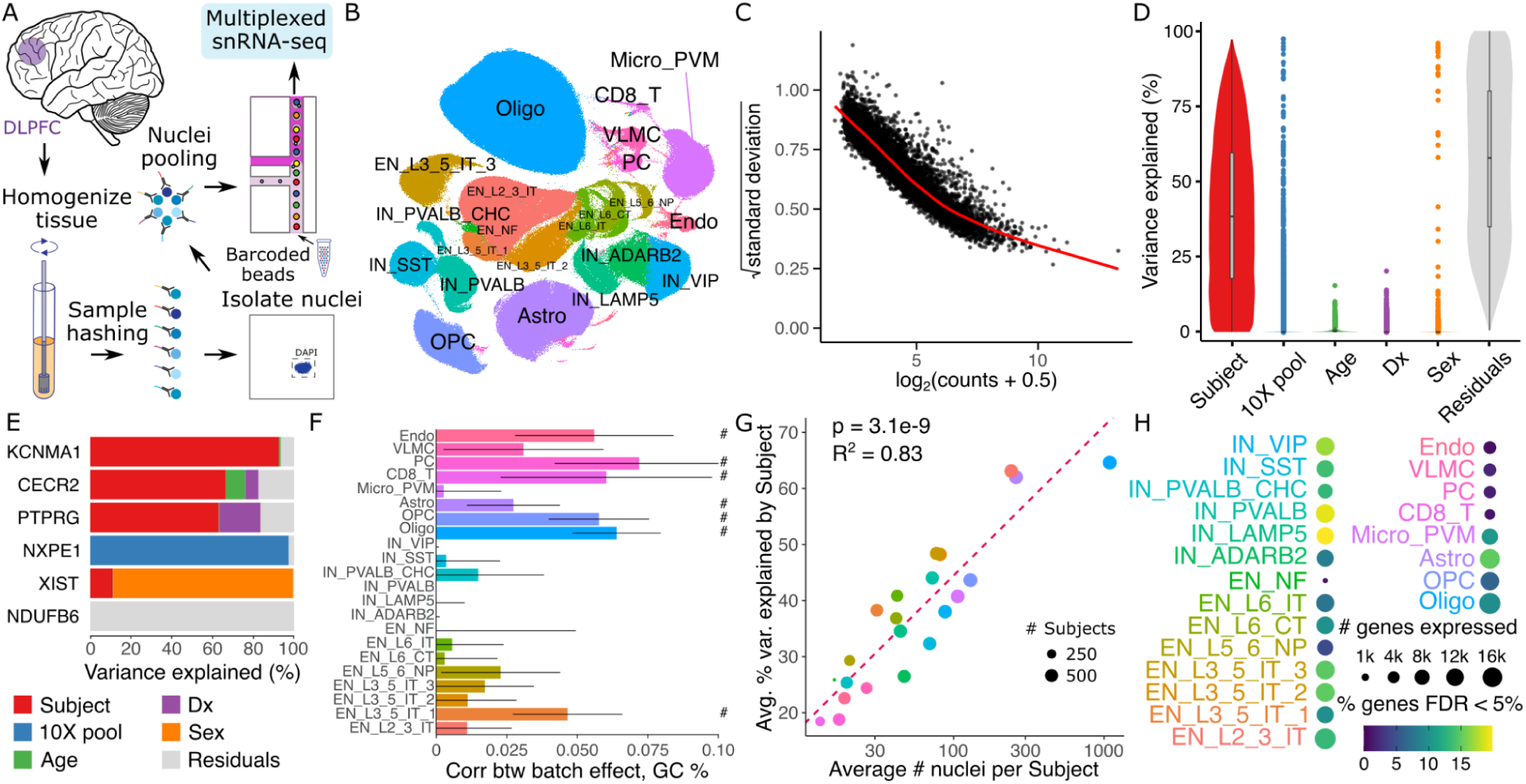
Dreamlet analysis of expression differences in Alzheimer’s Disease. **A)** Multiplexed single nucleus RNA-seq was performed on postmortem brain samples. **B)** UMAP dimensionality reduction with cell cluster annotations. **C)** Plot of mean-variance trend for expression counts in the microglia population. **D)** Violin plot for microglia summarizing variance partitioning analysis separating the fraction of expression variation for each gene into 6 components. **E)** Representative genes with high variance fractions explained by each of the 6 components in microglia. **F)** Spearman correlation of variance explained by sample pool ID with GC content of each gene. Bars indicate 95% confidence interval, ‘#’ indicates < 5% FDR. **G)** Annotated clusters with more nuclei per subject show higher concordance in technical replicates. Circle size indicates the number of subjects with at least 10 nuclei observed for the cluster. P-value from linear regression is shown. **H)** Number of genes passing expression cutoffs and the fraction of genes differentially expressed between AD subjects and controls at 5% FDR. Cell type abbreviations are: endothelial cells (Endo), vascular leptomeningeal cell (VLMC), pericyte (PC), CD8+ T-cells (CD8_T), microglia and perivascular macrophages (Micro_PVM), astrocytes (Astro), oligodendrocyte precursor cells (OCP), oligodendrocyte (Olig), inhibitory neurons (IN) and excitatory neurons (EN). IN annotations are followed by a marker gene, and EN are followed by a subtype annotation.

In microglia, the median fraction of variation explained by subject is 38.4%, indicating good reproducibility in expression values across technical replicates. The effect of AD status is modest with only 101 genes having more than 5% variance explained, underscoring the need for large sample sizes in order to characterize expression changes associated with the disease. While the sample pool explains minimal variance for the majority of genes, 932 genes have >5% variance explained across the pools. This indicates a substantial batch effect for these genes. The high dimensionality of the batch effect due to multiplexing 6 samples per pool indicates that modeling this as a random effect using dreamlet can retain high power while avoiding false positive findings. These findings are consistent across other cell types (**Supplementary** Figure 7). Further characterizing the batch effect reveals significant correlations between the variance across sample pools and the GC content of each gene (computed using all exons in the reference genome) in many cell clusters. This is consistent with previous work showing PCR artifacts driving technical variation ^36,37^ (**Figure 5F**).

While single cell/nucleus assays are uniquely capable of identifying cell type specific effects, the statistical power for each cell cluster in a given dataset varies widely. In particular, increasing the average number of nuclei per subject is directly related to an increase in the technical reproducibility of the gene expression measurements according to the variance explained across technical replicates from the same subject (**Figure 5G**). Moreover, the number of nuclei per subject impacts the read count per subject, the number of genes that pass a minimum expression cutoff, and the number of genes that are found to be differentially expressed between AD and controls (**Figure 5H**).

### Large effect up-regulation of *PTPRG* in microglia of Alzheimer’s Disease cases

The number of expressed genes and differentially expressed genes between donors with AD and controls varies widely between cell types and increases with the number of nuclei observed per subject (**Supplementary** Figure 8). The role of microglia in AD has received much recent attention due to findings from human genetic studies and mouse models ^38–40^, but the understanding of disease-associated gene expression changes has been more limited. Here, microglia have 1037 differentially expressed genes passing the study-wide 5% FDR cutoff (**Figure 6A**). Even so, *PTPRG* stands out with a log_2_ fold change of 1.59 and p-value of 2.94e-28. Genes demonstrate a range of cell type specificity patterns with many differentially expressed genes shared across multiple excitatory and inhibitory neuron subtypes (**Figure 6B**).

**Figure 6:**
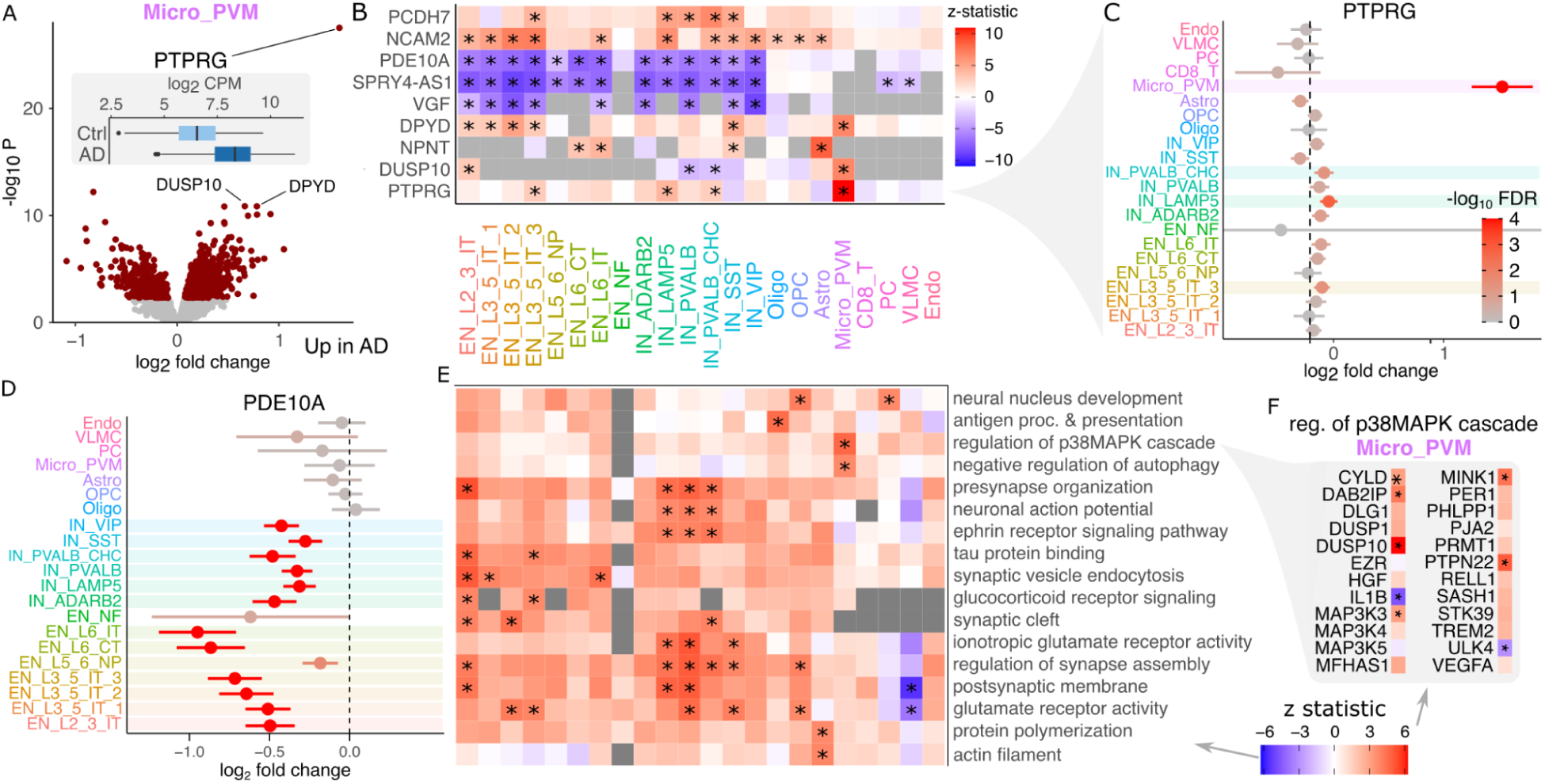
Gene expression signatures of Alzheimer’s disease. **A)** Volcano plot of differential expression between AD cases and controls in microglia. Red points indicate FDR < 5%. Inset shows boxplot of PTPRG stratified by disease status. **B)** Heatmap showing differential expression z-statistic for genes in each cell cluster. ‘*’ indicates study-wide FDR < 5% in all panels. Grey box indicates a gene did not pass expression cutoff in that cell cluster. **C)** Forest plot of log_2_ fold change for *PTPRG* in each cell type. Bars indicate 95% confidence interval. Color indicates FDR. **D)** Forest plot for *PDE10A*. **E)** Gene set analysis using the full spectrum of differential expression test statistics. **F)** Differential expression results for genes involved in regulation of p38MAPK cascade.

Notably, the *PTPRG* up-regulation with this large effect size is specific to microglia, with significant but much smaller log_2_ fold changes observed in 1 excitatory and 2 inhibitory neuron subtypes (**Figure 6C**). Other genes show effects across a range of cell types, as *PDE10A* is significantly down-regulated in AD in 13 of 14 neuron subtypes (**Figure 6D**). Gene set analysis using the full spectrum of test statistics in each cell cluster identifies up-regulation of synapse assembly and glutamate pathways in subsets of excitatory and inhibitory neurons, up-regulation of presynapse organization and synaptic vesicle endocytosis most strongly in EN_L2_3_IT, neuronal action potential most strongly in IN_PVALB, IN_PVALB_CHC and IN_LAMP5 (**Figure 6E**). In astrocytes there is a specific upregulation of protein polymerization. Oligodendrocyte precursors (OPC) have a specific upregulation of neural nucleus development. Notably, microglia have a specific up-regulation of the p38MAPK cascade, which is involved in microglial inflammatory response ^41^. This pathway includes *DUSP10* which is one of the top upregulated genes in microglia (**Figure 6F**).

## Discussion

The advent of single cell technology has enabled generation of large-scale high-resolution atlases to characterize cell type specific biology ^42–44^. As the scale of datasets continues to increase, there is new potential to study variation across subjects at the cell type level and how gene expression changes relate to a subject’s age, sex, disease state, and many other traits^6–10^.

We present the dreamlet software, an open-source R package that enables analysis of massive-scale single cell/nucleus transcriptome datasets. Dreamlet addresses both CPU and memory usage limitations by performing preprocessing and statistical analysis in parallel on multicore machines, and can distribute work across multiple nodes on a compute cluster. Dreamlet also uses the H5AD format for on-disk data storage to enable data processing in smaller chunks to dramatically reduce memory usage ^45^. The dreamlet workflow easily integrates into the Bioconductor ecosystem ^46^, and uses the SingleCellExperiment class ^47^ to facilitate compatibility with other analyses. Fitting precision-weighted linear mixed models ^23^ enables control of the false positive rate while retaining high power, even in the presence of high dimensional batch effects. Beyond differential expression testing, dreamlet provides seamless integration of downstream analysis including quantifying sources of expression variation ^48^, gene set analysis using the full spectrum of gene-level t-statistics ^33^ and visualizing results.

We also introduce a novel dataset of 1.4M single nuclei from postmortem brains from 150 Alzheimer’s disease cases and 149 controls. Analysis using the dreamlet software examines the cell type specific biology of AD. Highlighting the role of microglia in the disease, we observed that *PTPRG*, a protein tyrosine phosphatase receptor, is upregulated in AD and stands out substantially from all other genes in terms of effect size and p-value. PTPRG is an inflammatory marker but its role in the molecular etiology of AD is unclear ^49,50^. Interestingly, *PTPRG* is among the three genes (the other two being *APOE* and *DYPD*) that are reliably upregulated in two out of three previous human microglia transcriptome studies ^51^. Recent genome-wide association studies of AD do not identify risk variants in the region of the gene, suggesting that *PTPRG* upregulation is reactive, and might behave differently depending on the stage and progression of AD ^51^. The AD expression signatures identified here show high concordance with recent single nucleus data from Mathys ^52^ reanalyzed here with the dreamlet workflow, with overexpression of *PTPRG* in microglia being the top finding in both studies (**Supplementary** Figure 9,10).

Yet our work also highlights challenges in single-cell and -nucleus studies. First, technical batch effects can be substantial for some genes, and downstream analysis must account for these effects in order to control the false positive rate. Second, the findings within each cell cluster can be driven by biology, but also by limitations in the precision of gene expression measurements. We observe that increasing the measurement precision by increasing the number of reads and nuclei increases the reproducibility across technical replicates, and increases the number of differentially expressed genes identified. This wide variation in measurement precision across cell clusters yields wide differences in power to detect differential signals even within the same dataset. A finding that a gene is differentially expressed in only one cell cluster may be driven by cell type specific biology of disease, but could also be due to lower power in the other cell clusters. While the prospect of understanding the cell type specific biology of disease motivates these large cohort-scale studies, we recommend caution in the interpretation of cell type specific findings.

The dreamlet workflow focuses on differential expression across subjects by testing for changes in mean expression. Dreamlet, like all pseudobulk approaches, has some limitations. Aggregating gene expression across cells within a sample does not consider expression variation *within* a sample. Therefore, aggregating loses information about multimodal expression distributions and gene expression outliers within a sample seen in cases of stimulus response ^53,54^, and continuous expression gradients seen in developmental trajectories and analyses of pseudotime and RNA velocity ^17^. Addressing such challenges requires single-cell-level analyses.

In conclusion, here we introduce dreamlet, an open-source R package (DiseaseNeurogenomics.github.io/dreamlet) that addresses previous challenges to perform efficient differential expression analyses across subjects in large-scale single-cell datasets. Dreamlet is substantially faster and uses less memory, supports complex statistical models and better controls the false positive rate compared to existing workflows, providing an important tool to address the need of expanding single cell/nucleus transcriptome datasets.

## Online Methods

### Efficient computation of pseudobulk using on-disk memory

The dreamlet package creates pseudobulk data from the raw read counts for each sample and cell cluster stored in a SingleCellExperiment object ^47^. SingleCellExperiment is Bioconductor’s core data class for single cell data, and data from other formats (i.e. Seurat ^55^) can easily be converted to it. Dreamlet handles SingleCellExperiment objects to support storing large single cell datasets, either in-memory or on-disk, in a way that is seamless for the end user.

For large-scale studies, loading the entire dataset into memory can be prohibitive. Storing 18K genes across 2M cells with double precision would require 288 Gb memory. If the dataset is sparse so that 80% of the entries are zero, loading the entire dataset as a `sparseMatrix` object requires 58 Gb of data. Yet, instead of loading the entire dataset into memory, the data can remain on-disk and be accessed using an approach that takes advantage of the H5AD file format built on top of the HDF5 format ^45^. The zellkonverter package ^56^ uses a DelayedArray backend to provide a seamless interface to an on-disk H5AD dataset through the interface of the SingleCellExperiment class. This enables any analysis designed for the SingleCellExperiment class to take advantage of on-disk access to large-scale datasets. This can dramatically reduce memory usage while still retaining high performance.

Creating pseudobulk from a large dataset involves summing reads across a set of cells for each gene. While on-disk access to H5AD avoids loading the entire dataset into memory at the same time, creating pseudobulk requires accessing each entry of the dataset. For large datasets, the amount of time required for this step can vary dramatically depending on implementation details. Following extensive experimentation using R, Rcpp and C++ code, we use the beachmat library ^57^ to summarize each block containing all genes and a subset of cells into pseudobulk. Using dreamlet to compute pseudobulk from real single cell data reduces both compute time and peak memory usage by more than an order of magnitude. For large datasets of hundreds of donors, only dreamlet is tractable without purchasing an expensive high memory machine.

### Performance comparison for computing pseudobulk

H5AD files were created from real single nucleus RNA-seq data generated here using nuclei annotated using 12 cell type clusters. Donors were sampled with replacement to evaluate performance on up to 1,000 donors. We compared `aggregateToPseudoBulk()` in our dreamlet R package, `aggregate.Matrix()` in the Matrix.utils R package, `aggregateData()` in the muscat R package (which uses `scuttle::summarizeAssayByGroup()` in the backend), and `pegasus.pseudobulk()` in the pegasus python library ^26^. Each method was run 10 times for each condition and the average memory usage and CPU time are shown. Performance was assessed on a compute cluster where each run had access to 144 Gb memory. For a fair comparison, results are shown using only 1 thread to compute the pseudobulk values. (Dreamlet can use more threads to achieve faster performance at the expense of using more memory.)

We note that peak memory usage of native R code can vary dramatically based on available system memory, and when R chooses to invoke garbage collection. Despite substantial fluctuations, especially for memory usage, these performance results are very robust across multiple runs and changes to internal parameters.

### Precision-weighted linear mixed models

In widely cited work on differential expression analysis of RNA-seq data, Law, et al ^24^ demonstrate feasibility of modeling measurement uncertainty in a count response by weighting by the precision (i.e. reciprocal of observation-level variance). Importantly, they show that approximating the log transformed counts using a weighted linear regression can outperform NB regression that models counts explicitly but suffers from poor hypothesis testing for finite sample sizes (since the null distribution of the test statistics relies on asymptotic theory). Our motivation for using weighted linear regression to model transformed counts follows that of Law, et al ^24^, but becomes more pressing with repeated measures and complex study designs. In previous work on bulk RNA-seq, we have demonstrated that precision weighted linear mixed models are computationally efficient and hypothesis tests have good finite-sample performance in retaining power while controlling the false positive rate ^23^. Using generalized linear mixed models (GLMM) such as a negative binomial mixed model, can be very computationally demanding, suffer from convergence issues on real data, and produce poorly calibrated p-values on finite samples.

While Law, et al ^24^ consider precision weights to model heteroskedasticity of bulk RNA-seq counts, here we consider two levels of precision weights.

### Modeling measurement uncertainty in count models using a two-stage weighting approach

The original biospecimen is often composed of thousands of cells of a particular type, or corresponding to a particular empirically defined cell cluster. In order to study the biology of a given cell type, experimental workflows randomly sample single cells for RNA sequencing. Since this is a stochastic process that samples cells from a much larger population, gene expression measurements aggregated across a larger number of cells more precisely represents expression in the full population of cells. Consequently, the precision of expression measurements of a gene across samples is directly related to the number of cells sequenced per specimen. Statistically, gene expression measurements vary in their precision and are thus heteroskedastic.

The limited read count per gene in RNA-seq experiments has a similar effect on measurement uncertainty, with more reads giving a more precise measurement on the log scale. The widely used limma-voom approach ^24^ estimates precision weights by fitting a linear model for each gene. Voom then smoothes the relationship between the log2 counts per million and the square root residual standard deviation from the model fit of each gene. We have previously extended the voom approach to enable fitting of linear mixed models for estimating precision weights ^23^. Here we further extend this work to allow modeling of heteroskedasticity using initial precision weights. The resulting estimated precision weights incorporate measurement uncertainty due to stochastic sampling effects at both the level of cells and reads.

Here, we initialize the precision weights that are fed into an empirical mean-variance trend fit by using an approximation to a Poisson generative model of the pseudobulk read counts. See **Supplementary Methods** for details.

### Empirical Bayes shrinkage for precision-weighted linear mixed models

For small sample sizes, parameter estimates can have high sampling variance. In seminal work, Smyth ^25^ developed an empirical Bayes approach that borrows information across genes to estimate the residual variance. The widely used limma package ^58^ fits a linear model for each gene, performs the empirical Bayes step, and then computes a moderated t-statistic with a modified null distribution. In the case of a linear model, Smyth’s empirical Bayes method uses a conjugate prior on the residual variances and assumes they are drawn from an inverse gamma distribution (i.e. precisions are drawn from a scaled chi-squared distribution) with parameters estimated from the data. A key value in this calculation is the residual degrees of freedom.

In the case of a linear model with *n* samples and *p* covariates (including the intercept), the residual degrees of freedom (*df_r_*) is simply *n - p*. However, the case of a linear mixed model used here is more complicated. In this case, we show that the residual variance estimates follow a distribution given by a weighted mixture of *n* chi-squared random variables, where the weights depend on both the data and the estimated model parameters (**Supplementary Information**). We match the expected value of this mixture distribution using a single chi-square and use its degrees of freedom to approximate the *df_r_* of the linear mixed model. Importantly, this method is exact in the case of a linear model, is approximate for linear mixed models with any number of random effects, and the approximation improves with the sample size.

### Analysis of simulated single cell data

We followed the workflow from Cromwell, et al. ^18^ to estimate parameters from read single cell data and then simulate data based on this. We added our dreamlet software to their existing workflow and used performance metrics as described in their paper. We also added code to compute the area under the precision recall curve (AUPR) and F1 scores to evaluate statistical performance. Finally, we extended the simulations to vary the number of cells observed for each sample in order to better match real data. Instead of specifying the exact number of cells observed per sample, the average number is specified and the number for each sample is drawn from a Dirichlet-multinomial distribution.

### Analysis of real single nucleus data with permuted disease labels

We used the single nucleus data generated here and randomly permuted the disease labels of each donor while retaining the same fraction of Alzheimer’s disease cases and controls within each sample batch. Each method was then run with default parameters.

### Single nucleus RNA-seq data generation

#### Study cohort

Frozen brain tissue samples derived from DLPFC (Brodmann area 9/46) were obtained from the Mount Sinai Brain Bank (MSBB–Mount Sinai NIH Neurobiobank), which holds over 2,000 human brains. Since we wanted to leverage samples from donors with either no discernable neuropathology or cognitive complaints (controls), or with only AD-associated neuropathology, we narrowed our initial selection of brain donors using a combination of neuropathological and clinical criteria inspired by previous work ^59,60^. AD samples needed to be classified by (1) CERAD protocol ^61^ as “AD possible”, “AD probable” or “AD definite”, (2) Braak AD staging protocol ^62^ within the stage 3-6, and (3) clinical dementia rating ^63^ within the rate 0.5-5.

Furthermore, our subset of AD donors cannot be diagnosed with Parkinson’s disease or Diffuse Lewy body disease. Conversely, control samples needed to be classified by (1) CERAD protocol as “no AD” or “possible AD” and (2) Braak AD staging protocol within the stage 0-2. Additionally, control samples cannot be diagnosed with any other neurodegenerative, neurological or neuropathological diagnosis. Neuropathological assessments, cognitive, medical status and neurological status were performed according to established procedures ^64^. All neuropsychological, diagnostic and autopsy protocols were approved by the Mount Sinai and JJ Peters VA Medical Center Institutional Review Boards.

#### Isolation and fluorescence-activated nuclear sorting (FANS) of nuclei with hashing

All buffers were supplemented with RNAse inhibitors (Takara). 6 samples were processed in parallel. 25mg of frozen postmortem human brain tissue from each specimen was homogenized in cold lysis buffer (0.32 M Sucrose, 5 mM CaCl_2_, 3 mM Magnesium acetate, 0.1 mM, EDTA, 10 mM Tris-HCl, pH8, 1 mM DTT, 0.1% Triton X-100) and filtered through a 40 µm cell strainer. The flow-through was underlaid with sucrose solution (1.8 M Sucrose, 3 mM Magnesium acetate, 1 mM DTT, 10 mM Tris-HCl, pH8) and centrifuged at 107,000 g for 1 hour at 4 °C. Pellets were resuspended in PBS supplemented with 0.5% bovine serum albumin (BSA). Resuspended nuclei were quantified (Countess II, Life Technologies) and 2M from each sample were pelleted at 500 g for 5 minutes at 4°C and re-suspended in 100 µl staining buffer (2% BSA, 0.02% Tween-20 in PBS). Each sample incubated with 1 µg of a distinct TotalSeq-A nuclear hashing antibody (Biolegend) for 30 min at 4 °C. Prior to FANS, volumes were brought up to 250 µl with PBS and 7-AAD (Invitrogen) added to facilitate detection of nuclei. 7-AAD positive nuclei were collected in tubes pre-coated with 5% BSA using a FACSAria flow cytometer (BD Biosciences).

#### snRNA-seq and library preparation

Following FANS, nuclei were washed twice in staining buffer before being re-suspended in 22 µl PBS and quantified. Nuclei concentrations were normalized and equal amounts from each sample were pooled together. 2 aliquots of 60,000 pooled nuclei (i.e. 10,000 each) were processed in parallel using 3’ v3.1 reagents (10x Genomics). At the cDNA amplification step (step 2.2), reactions were supplemented with a hash-tag oligo (HTO) cDNA “additive” primer (GTGACTGGAGTTCAGACGTGTGCTCTTCCGAT*C*T; *Phosphorothioate bond). Following cDNA amplification, supernatants from the 0.6x SPRI (Beckman Coulter) selection step were retained for HTO library generation. Otherwise, cDNA libraries were prepared according to the manufacturer’s instructions (10x Genomics). HTO libraries were prepared as described previously ^65^. All libraries were sequenced at NYGC using the Novaseq platform (Illumina).

### Single nucleus data processing and quality control

#### Alignment and demultiplexing

Sequencing reads from all pools of multiplexed samples were aligned to the hg38 reference genome using STARsolo ^66,67^. To assign the cells from each pool to their respective donors, we applied a genotype-based demultiplexing approach followed by genotype concordance. First, cellSNP ^68^ was used to pile up the expressed alleles from polymorphic sites overlapping snRNA-seq reads. Then, vireo ^13^ utilized those pile-ups to split cells into clusters corresponding to six distinct donors per pool. The assignment of identity of each cluster of cells to a particular donor was derived from genotype concordance analysis that compared the clusters of cells against reference SNP-array data using QTLtools-mbv ^69^. While the majority of pools contained the cells from the expected sets of donors, we leveraged the genotype concordance results to detect and correct occasional sample swaps and mislabelings.

### Quality control, UMAP, cell type annotation

**QC.** After genome alignment and demultiplexing, we applied rigorous three-step QC to remove ambient RNA and retain viable nuclei for downstream analysis. First, the QC was applied at the cell level. A battery of QC tests was performed to filter low-quality libraries and non-viable cells within each library. Poor-quality cells were detected by thresholding based on UMI counts, gene counts, and mitochondrial contents. We also checked for possible contamination from ambient RNA, a fraction of reads mapped to non-mRNA like rRNA, sRNA, and pseudogenes, as well as known confounding features such as lncRNA *MALAT1*. Further filtering was carried out by removing doublets using the Scrublet method^70^. Second, the QC is applied at the feature level. We removed features (genes) that are not robustly expressed by at least 0.05% of the cells/nuclei. Lastly, the QC was applied at the donor level. We remove donors with less than 50 cell counts, which can introduce more noise to the downstream analysis.

#### Batch correction

We have developed a tracking platform to record all technical covariates (such as 10x Genomics lot kit number, dates of different preparations, viable cell counts, etc.) and quality metrics derived from data preprocessing. We assessed the correlation between all pairs of technical variables using Canonical Correlation Analysis and used the Harmony method^71^ to regress out the effect of sequencing pools before performing clustering and taxonomy analysis.

#### Clustering

Highly variable features were selected from mean and variance trends, and we used the k-Nearest-Neighbor (kNN) graph calculated on the basis of harmony-corrected PCA embedding space to cluster cells in the same cell-type using Leiden^72^ clustering algorithm. We used UMAP ^73^ for the visualization of resulting clusters.

#### Cellular taxonomy

Identified cell types will be annotated based on a combination of expert curation and machine-learning-based algorithms to query known gene marker signatures previously curated by Human Cell Atlas.

#### Sex check

For each donor the labeled sex was checked to be consistent with expression on *XIST* and *UTY* genes on the X and Y chromosomes, respectively.

## Code Availability

The dreamlet R package, including documentation, tutorials and code examples, is available at DiseaseNeuroGenomics.github.io/dreamlet and is available on Bioconductor at https://bioconductor.org/packages/dreamlet/. Code for simulations is available at github.com/GabrielHoffman/muscat-comparison_v3. Data analysis code and results for analyses in Figures 3-6 are available at https://github.com/GabrielHoffman/dreamlet_analysis. The snRNA-seq data generated here is available at https://www.synapse.org/PsychAD_public.

## Software versions

dreamlet v1.1.24,

muscat v1.11.2,

DESeq2 v1.36.0,

limma 3.52.1,

edgeR 3.38.0,

MAST 1.22.0,

pegasus 1.6.0

STARsolo 2.7.9.

cellSNP 1.2.0

vireo 0.5.6

QTLtools-mbv 1.3

## Author Contributions

G.E.H. developed the dreamlet package and performed analysis. V.H. provided tissue specimens from Alzheimer’s disease brains and controls. A.H., C.C., M.A., Z.S. and S.A. generated novel snRNA-seq data from Alzheimer’s disease brains and controls under the supervision of J.F.F. D.L., J.B., P.F., K.T., S.V., G.V. performed processing of snRNA-seq data from Alzheimer’s disease brains and controls generated here. G.E.H. and P.R. supervised analysis. G.E.H., D.L., J.B., J.F.F. and P.R. wrote the manuscript with input from all authors.

