## Supplement for "Efficient differential expression analysis of large-scale single cell transcriptomics data using dreamlet"

### 1 Supplementary Figures

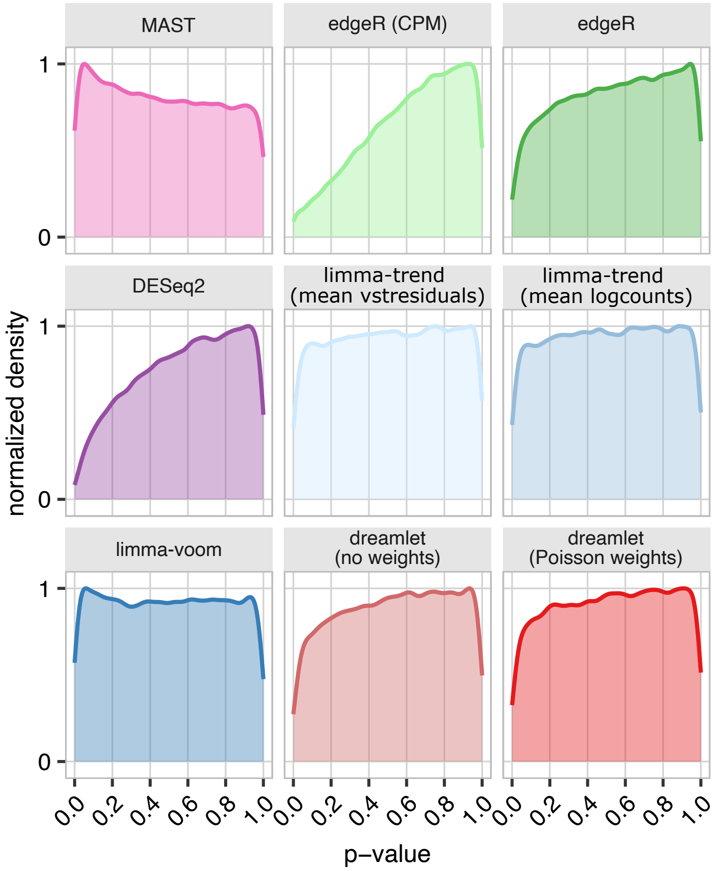

**Supplemental Figure 1: Distribution of p-values under the null model.** Density plot of p-values from 4000 genes, 20 samples and 200 cells per sample under a null model where there is no true association between gene expression and the variable tested.

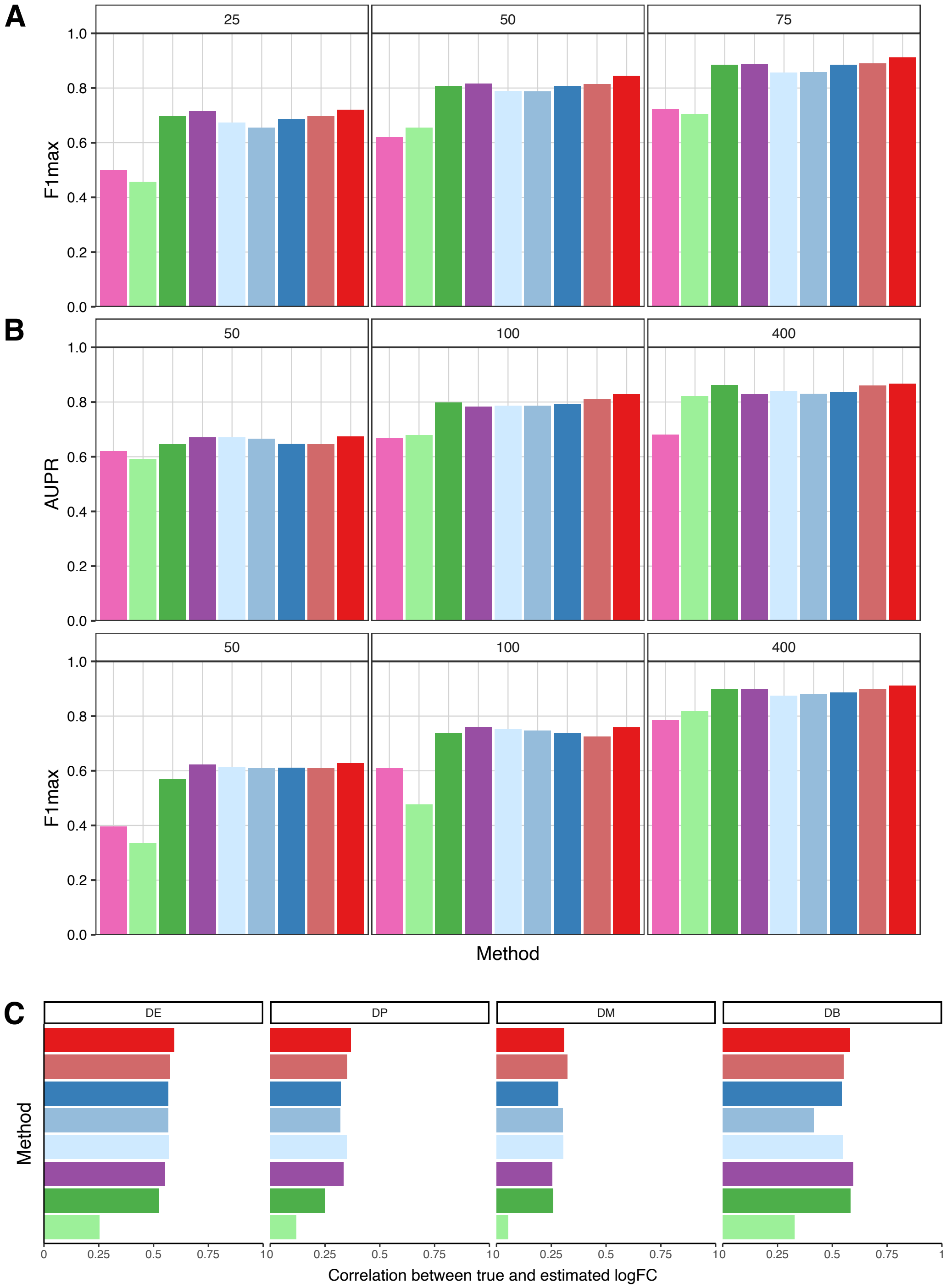

**Supplemental Figure 2: Performance metrics for differential expression methods. A)** Performance with sample size increasing from 25 to 75 with a mean of 400 cells per sample measured by maximum F1 score. This is based on the same simulations as in Figure 2D. **B)** Performance with the number of cells per sample increasing from 50 to 400 for 100 samples measured by AUPR and maximum F1 score. **C)** Pearson correlation between true and estimated log fold change coefficients. Results are shown for 4 types of differential expression: change in mean expression (DE), change in proportions of high and low expression (DP), differential modality (DM) and both a change in proportions and modality (DB). See Crowell et al. (2020) for details of simulation models.

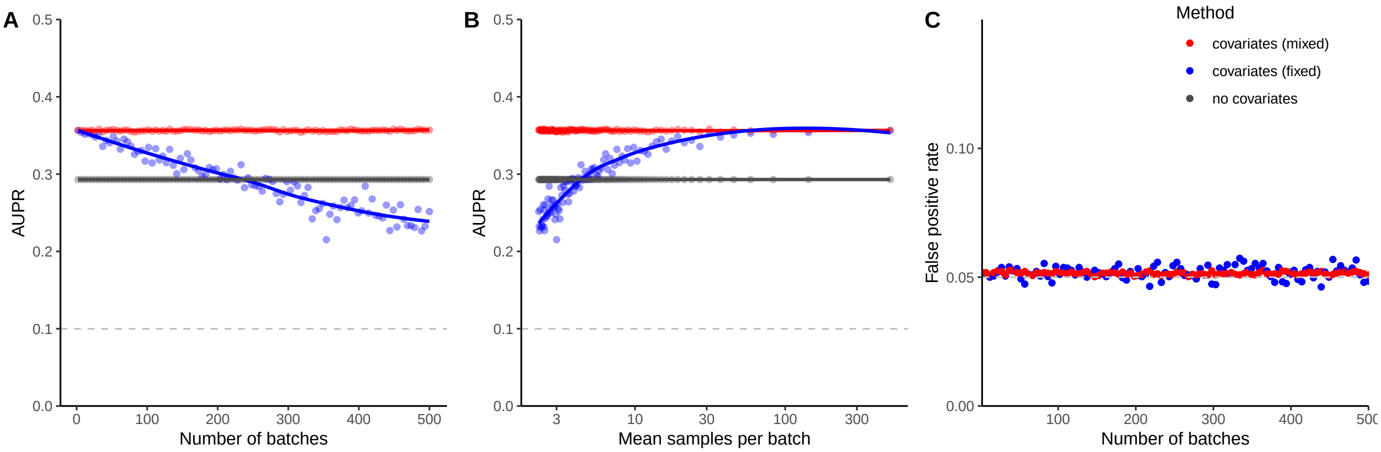

**Supplemental Figure 3: Performance of mixed models on simulated data**. Simulations were performed for $n = 1000$ samples, and an increasing number of batches where the effect of each batch is drawn from a standard normal distribution. For each number of batches, 5000 genes were simulated and 500 were associated with a phenotype variable that was normally distributed. For each simulation, regression models accounting for batch using a fixed or mixed effect model, or no covariate correction were evaluated. The area under the precision recall curve (AUPR) and false positive rate (FPR) at 5% was computed for each simulation, and a lowess smooth is plotted. **A**) AUPR for each model versus the number of batches in the simulation. **B**). The same results as in **(A)** shown as a function of mean number of samples per batch on a log scale. **C**) FPR as a function of the number of batches. Both fixed and mixed effects models control the false positive rate, but the mixed model shows substantially reduced variance around the 5% target compared to the fixed effect model. Omitting covariates give inflated FPR beyond the y-axis shown here.

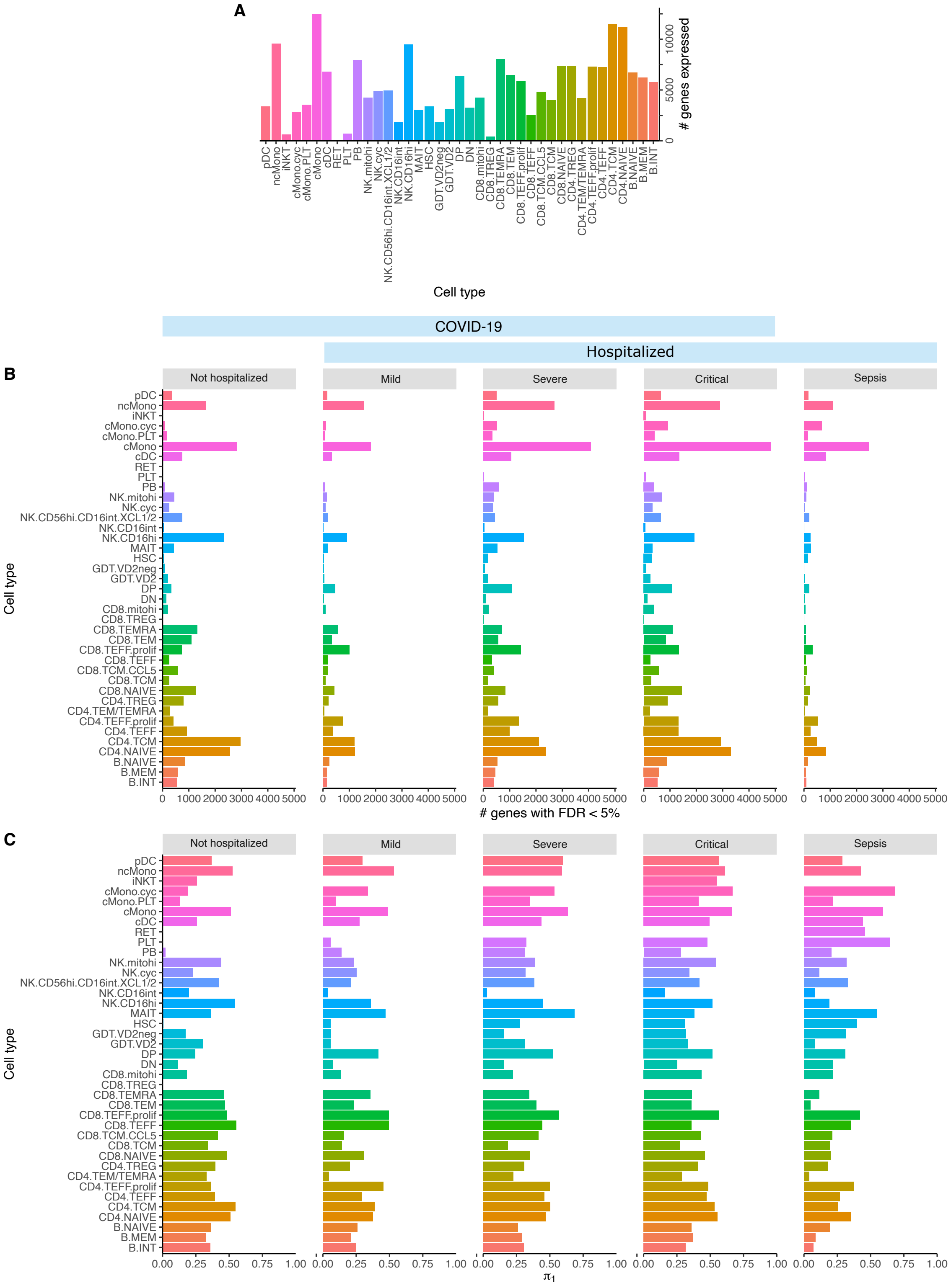

**Supplemental Figure 4: Summary of differential expression results for COVID-19 severity. A)** Number of genes passing expression cutoff for each cell type. **B)** Number of differentially expressed genes at FDR 5%. for each disease stated compared to healthy controls. **C)** Storey's $\pi_1$ estimate of the fraction of genes rejecting the null hypothesis of no association.

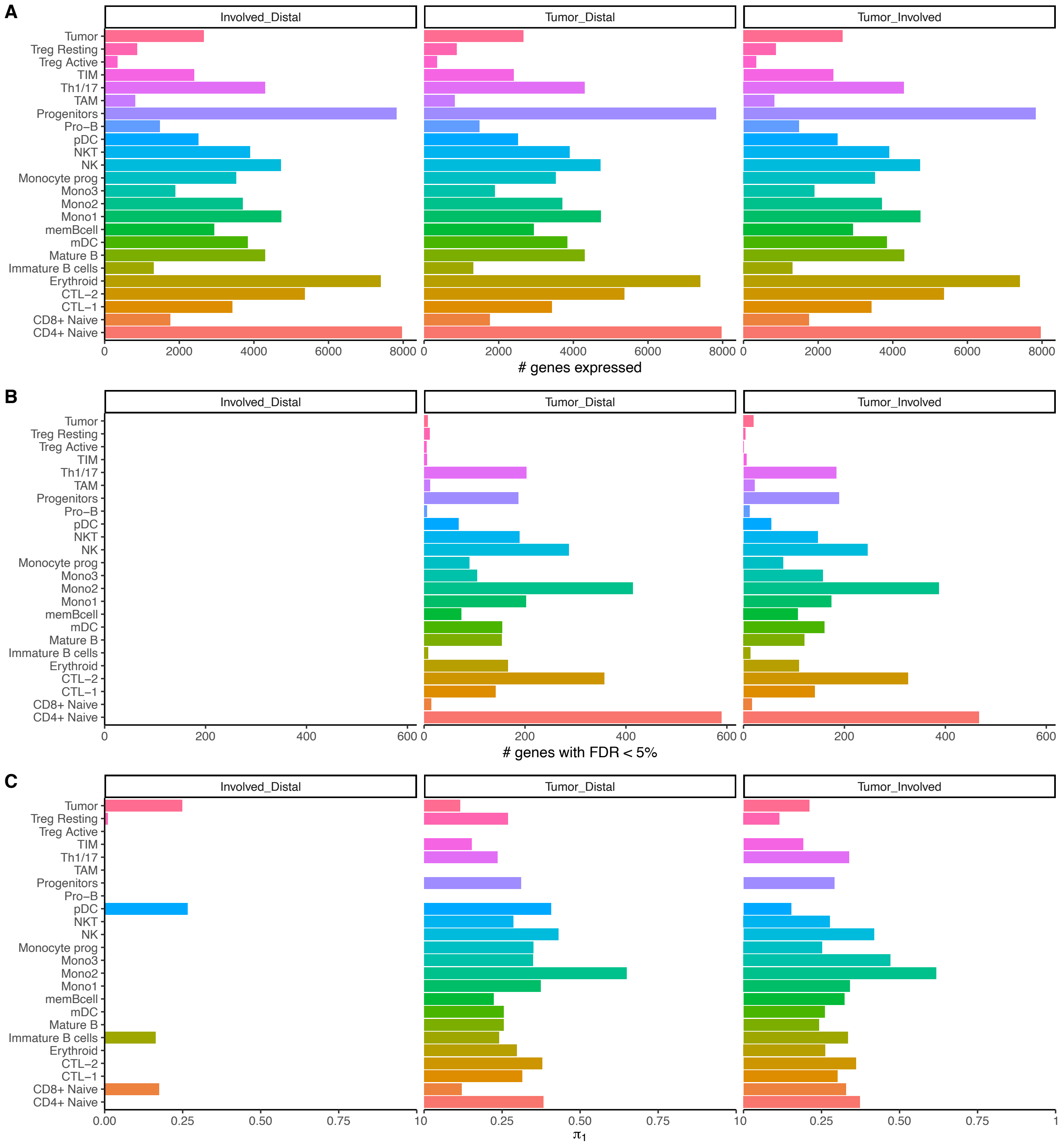

**Supplemental Figure 5: Summary of differential expression results bone cancer metastases . A)** Number of genes passing expression cutoff for comparison between tumor, involved tissue and distal tissue for each cell type. **B)** Number of differentially expressed genes at FDR 5%. Note that no genes passed the cutoff for the "Involved vs Distal" comparison. **C)** Storey's $\pi_1$ estimate of the fraction of genes rejecting the null hypothesis of no association.

**A**

| | Female | Male |
|---|---|---|
| Control | 70 | 79 |
| AD | 71 | 79 |

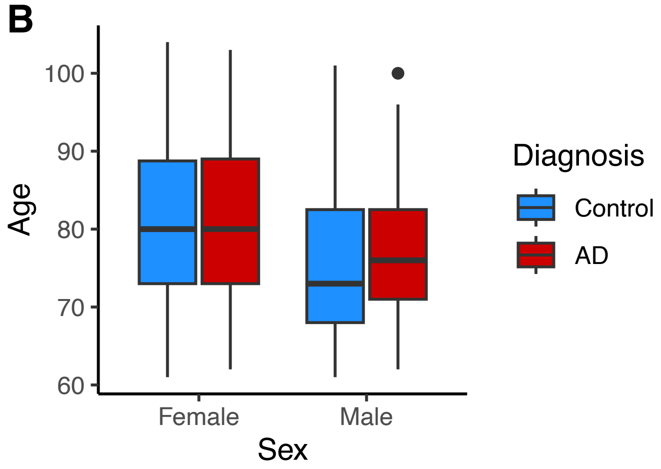

**Supplemental Figure 6: Summary of Alzheimer's disease postmortem brain cohort A)** Sample size stratified by sex and disease status **B)** Age distribution stratified by sex and disease status.

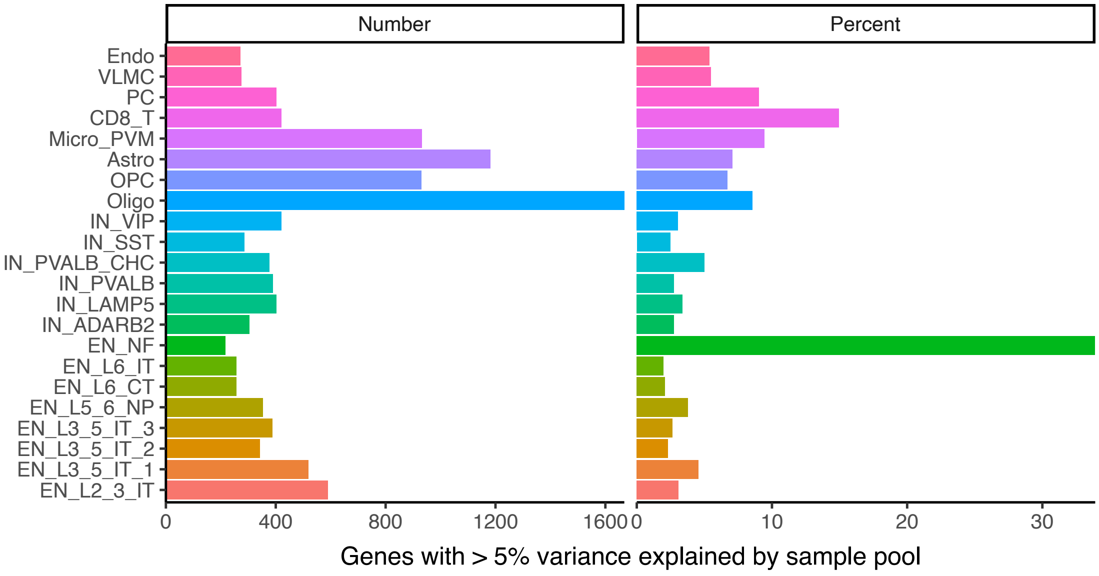

**Supplemental Figure 7: Summary of batch effect across sample pools** The number (left) and percentage (right) of expressed genes in each cell type where > 5% of variance is explained by variation across sample pools.

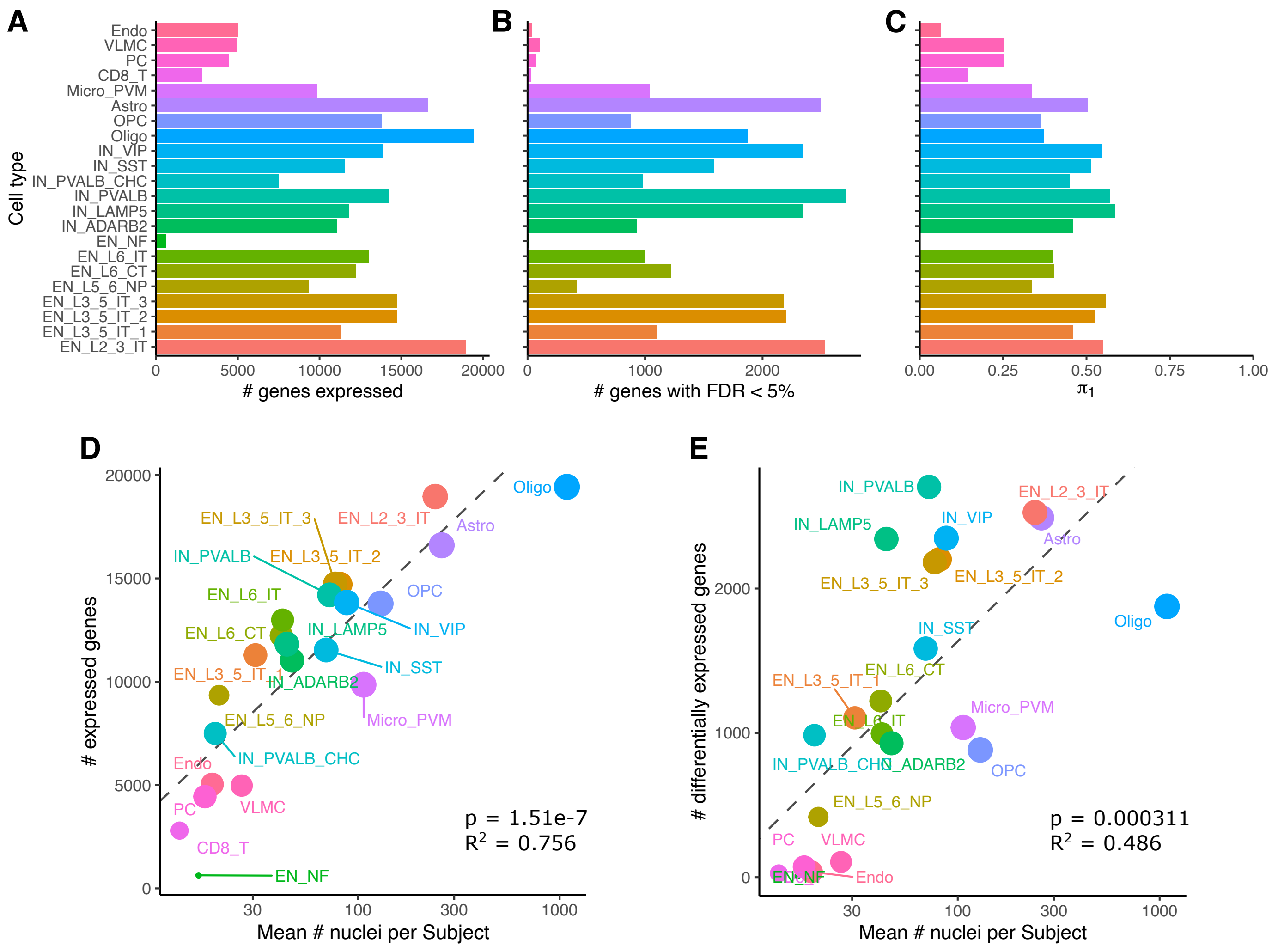

**Supplemental Figure 8: Summary of Alzheimer's disease differential expression analysis**. **A)** Number of genes passing expression cutoff for each cell type. **B)** Number of differentially expressed genes at 5% FDR. **C)** Storey's $\pi_1$ estimate of the fraction of genes rejecting the null hypothesis of no association. **D,E)** Number of (**D**) genes passing expression cutoffs and (**E**) differentially expressed genes for each cell cluster increases with the number of nuclei observed per subject. Circle size indicates the number of subjects with at least 10 nuclei observed for the cluster. P-value and R-squared from linear regression is shown.

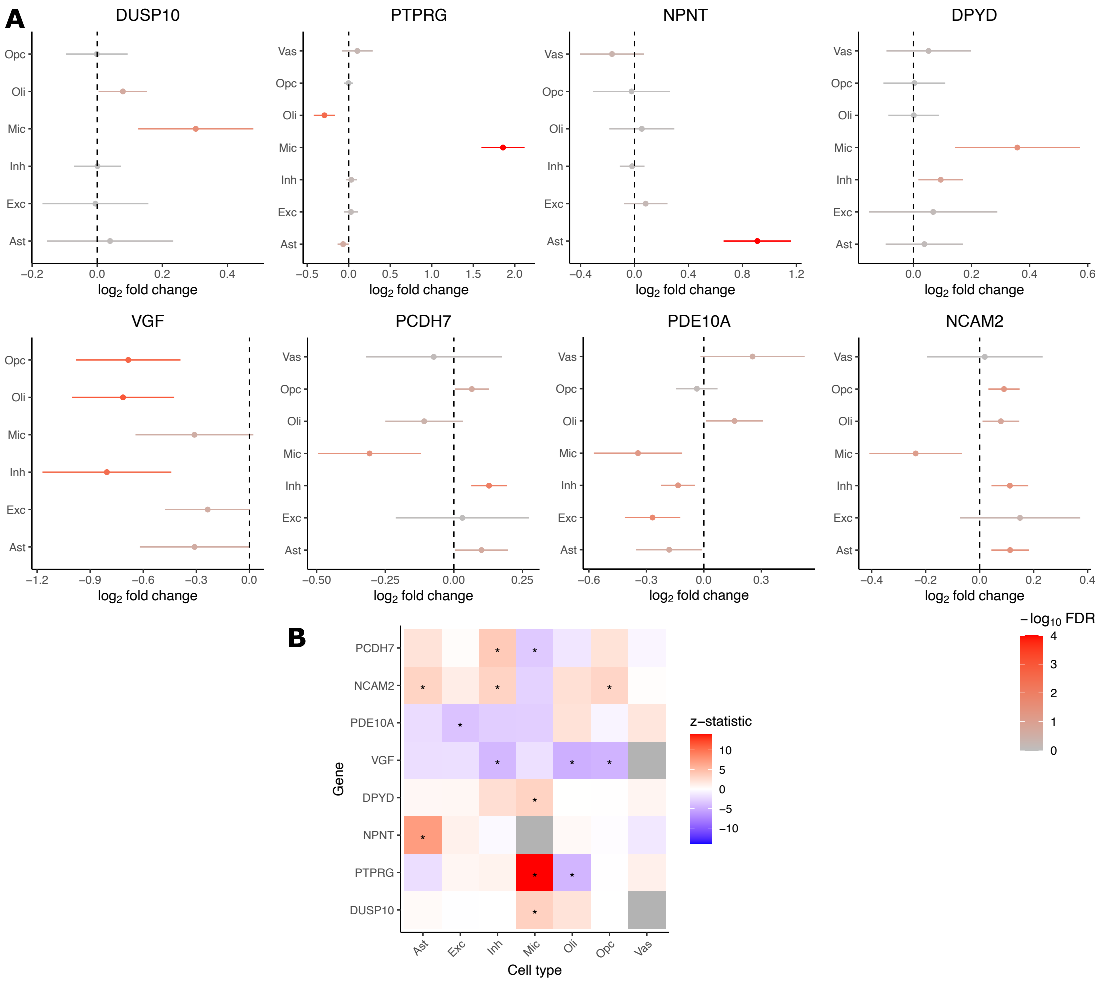

**Supplemental Figure 9: Replication of cell type patterns in Alzheimer's disease using dreamlet analysis.** Examples of logFC from Mathys et al. (2023) reproduce cell type patterns from current work. Genes highlighted here were selected based on analysis of PsychAD data in the main text. **A)** Forest plot of logFC and 95% confidence intervals. Color indicates FDR. **C)** Heatmap indicating z-statistic for differential expression between Alzheimer's disease and controls. Cell annotations from Mathys et al. (2023) are excitatory neurons (Exc), inhibitory neurons (Inh), oligodendrocytes (Oli), oligodendrocyte precursor cells (OPC), astrocytes (Ast), microglia (Mic), and vascular cells (Vas).

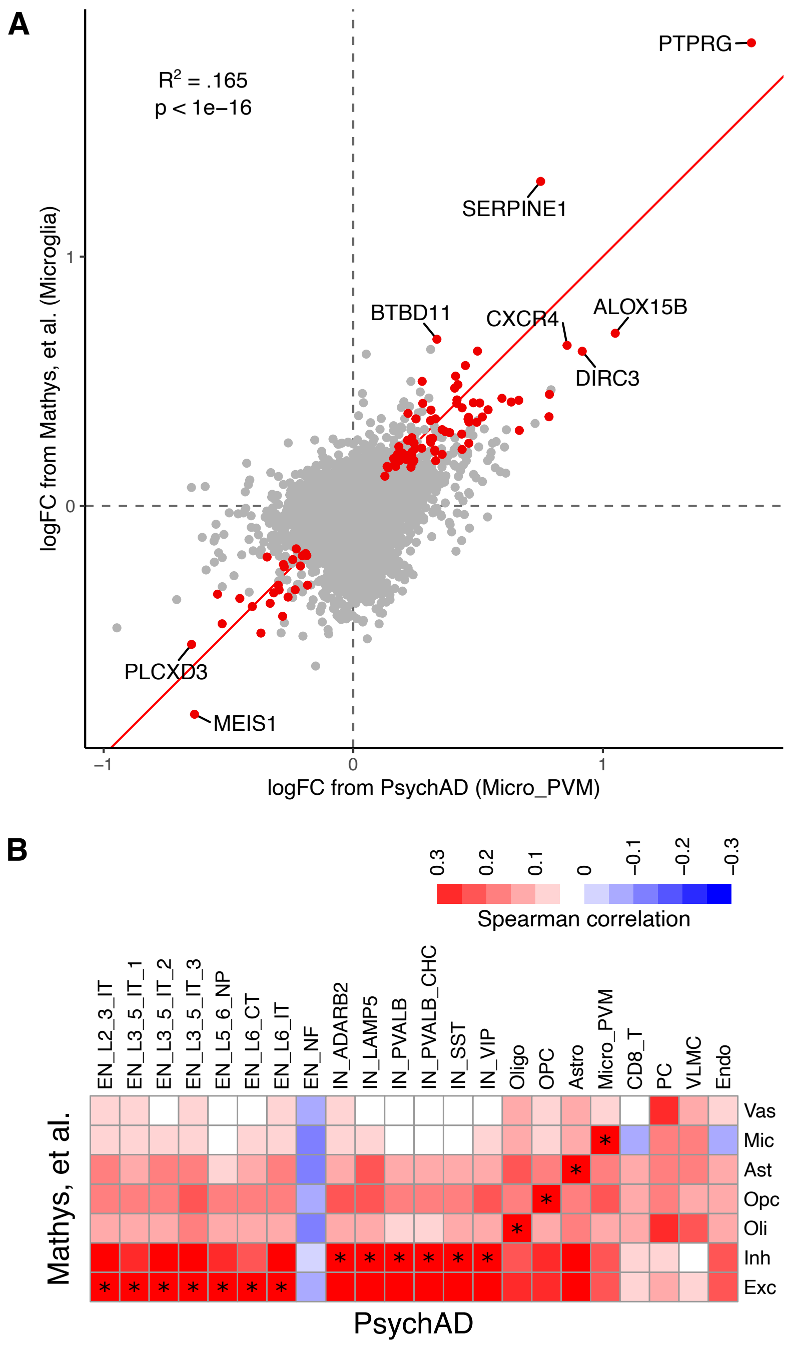

**Supplemental Figure 10: Genome-wide replication in effect size for Alzheimer's disease differential expression analysis**. Concordance of log fold change (logFC) from current work compared to Mathys et al. (2023) **A)** Plot of logFC for Alzheimer's disease versus controls for both studies in microglia. Red points indicate genes that are study-wide significant in both datasets. Red line indicates equal logFC. Spearman correlation is shown. **B)** Spearman correlation between logFC from cell types annotated in each study. '*' indicates match in cell type between studies based on annotation label. Note that the current work included multiple subtypes of excitatory and inhibitory neurons compared to single 'excitatory' and 'inhibitory' categories from Mathys et al. (2023). Cell annotations from Mathys et al. (2023) are excitatory neurons (Exc), inhibitory neurons (Inh), oligodendrocytes (Oli), oligodendrocyte precursor cells (OPC), astrocytes (Ast), microglia (Mic), and vascular cells (Vas).

### 2 Supplementary Methods

#### 2.1 Implementation of dreamlet workflow

The `dreamlet` workflow depends heavily on our `variancePartition` R package (Hoffman and Roussos, 2021, Hoffman and Schadt, 2016). The technical contributions of two-stage precision weights and empirical Bayes shrinkage for linear mixed models described in this manuscript are implemented in the `variancePartition` R package. The `dreamlet` package is a frontend designed to integrate with the Bioconductor ecosystem for single cell data, visualize results, and give users an interface to apply the updated statistical modeling implemented in `variancePartition`.

**Two-stage precision weights**
Standard `limma::voom()` estimates the precision weights by 1) fitting a regression model for each gene, 2) computing the residuals, 3) fitting a non-parametric mean-variance trend to the residuals, and 4) using this trend to compute the precision weights. This approach is adopted by our `dream` pipeline using `variancePartition::voomWithDreamWeights()` which performs step (1) allowing a linear mixed model instead of a fixed effects model that limma uses (Hoffman and Roussos, 2021).

Here we extend `variancePartition::voomWithDreamWeights()` to use initial precision weights at the observation-level in the regression models fit in step (1). By modeling uncertainty at the observation-level, each gene for each sample has a corresponding initial precision weight. This follows work by the developers of the `limma` package who developed 'quality weights' to downweight outliers in bulk RNA-seq data (Liu et al. 2015). That approach weights each sample while using the same sample weight across all genes. It does not model observation-level uncertainty and is not easily extended to the case of linear mixed models that we use here.

In addition, we improve the loess smoothing of the empirical mean-variance trend. The `limma::voom()` function and previous versions of `variancePartition::voomWithDreamWeights()` used a tuning parameter (i.e. `span`) that is set to a value of 0.5, although it can be manually changed by the user. Instead of using a fixed value or having a user change it by examining diagnostic plots, we apply an automated search of the parameter space and select a value based on an objective metric of the model fit. This is implemented by `fANCOVA::loess.as()` using a bias-corrected AIC criterion (Hurvich et al., 1998) and is accessed using `variancePartition::voomWithDreamWeights(...,span='auto')`.

**Empirical Bayes shrinkage for moderated t-statistics**
We extend the empirical Bayes approach of `limma` (Smyth, 2004) to the case of linear mixed models. The function new `variancePartition::eBayes()` takes the output of the `dream()` and performs empirical Bayes shrinkage to both linear and linear mixed models. From the user's perspective, it works just like `limma::eBayes()`. The standard workflow runs `variancePartition::dream()` to fit regression models, `variancePartition::eBayes()` to perform shinkage, and `variancePartition::topTable()` to extract results. The empirical Bayes step is not required,

and in that case `variancePartition::topTable()` will return results without applying moderated t-statistics.

#### 2.2 Initial precision weights

##### 2.2.1 A Poisson generative model of read counts

Consider cell $j$ from sample $i$ having observed count $c_{i,j}^{(g)}$ for gene $g$ and total read count $l_{i,j} = \sum_g c_{i,j}^{(g)}$. Hereafter, we suppress the $g$ term for simplicity since we only consider a single gene at a time. Let the observed count for a given gene be Poisson distributed according to $c_{i,j} \sim Pois(p_{i,j}\, l_{i,j})$ where $p_{i,j}$ is the expression fraction corresponding to this gene. Consider computing pseudobulk counts as $\tilde{c}_i = \sum_{j=1}^{n_i} c_{i,j}$ by summing reads across $n_i$ cells from sample $i$, and let the average library size from these cells be $\bar{l}_i = \frac{1}{n_i}\sum_{j=1}^{n_i} l_{i,j}$. Based on summing independent Poisson random variables, it follows that

$$
\begin{aligned}
\tilde{c}_i &\sim Pois\left(\sum_{j=1}^{n_i} p_{i,j} l_{i,j}\right) && (1)\\
&\sim Pois\left(n_i \bar{p}_i \bar{l}_i\right), && (2)
\end{aligned}
$$

where $\bar{p}_i$ is the mean expression fraction across the cells in sample $i$.

Let the counts per million (CPM) for pseudobulk computed from sample $i$ be $a_i = 1e6 \cdot \tilde{c}_i/(n_i \bar{l}_i)$. Based on properties of the Poisson distribution the mean and variance of $a_i$ are

$$
\begin{aligned}
E\left[a_i\right] &= E\left[1e6 \cdot \tilde{c}_i/(n_i \bar{l}_i)\right] && (3)\\
&= 1e6 \cdot E\left[\tilde{c}_i\right]/(n_i \bar{l}_i) && (4)\\
&= 1e6 \cdot (n_i \bar{p}_i \bar{l}_i)/(n_i \bar{l}_i) && (5)\\
&= 1e6 \cdot \bar{p}_i && (6)\\
\text{var}\left[a_i\right] &= \text{var}\left[1e6 \cdot \tilde{c}_i/(n_i \bar{l}_i)\right] && (7)\\
&= 1e12 \cdot \text{var}\left[\tilde{c}_i\right]/(n_i \bar{l}_i)^2 && (8)\\
&= 1e12 \cdot (n_i \bar{p}_i \bar{l}_i)/(n_i \bar{l}_i)^2 && (9)\\
&= 1e12 \cdot \bar{p}_i/(n_i \bar{l}_i) && (10)
\end{aligned}
$$

Now consider the variance of the log CPM. (Note that we use the natural logarithm here for simplicity, but processing of real data uses base 2 logarithm. Since converting between logarithm with different bases involves scaling by a constant factor, results here are proportional to results using a different base.)

Using the delta method to approximate transformations of random variables gives

$$\begin{aligned}
E\left[\log(a_i)\right] &\approx \log(E\left[a_i\right]) & (11)\\
&= \log(1e6) + \log(\bar{p}_i) & (12)\\
\mathrm{var}\left[\log(a_i)\right] &\approx \mathrm{var}(a_i)/E\left[a_i\right]^2 & (13)\\
&= \frac{1e12 \cdot \bar{p}_i/(n_i\bar{l}_i)}{(1e6 \cdot p)^2} & (14)\\
&= \frac{1}{n_i\bar{p}_i\bar{l}_i} & (15)
\end{aligned}$$

The formula based on the generative model recapitulates the intuition that increasing expression rate, number of cells and mean library size all reduce variation in log CPM. While $n_i$ and $\bar{l}_i$ are observed values, the variance depends on the unknown parameter $\bar{p}_i$. In real data, $n_i\bar{p}_i\bar{l}_i$ is the expected number of counts, so the variance can be estimated as $\widehat{\mathrm{var}}\left[\log(a_i)\right] = 1/\tilde{c}_i$. Since the precision is the inverse of the variance, the estimated precision of the log CPM is the observed number of counts:

$$\boxed{\widehat{\mathrm{prec}}\left[\log(a_i)\right] = \tilde{c}_i} \quad (16)$$

**Integration with voom-style precision weights**
The widely used voom method (Law et al., 2014) fits an initial unweighted regression model to each gene, fits a smooth curve to the empirical mean-variance trend where the variance values are computed from the residual variances from the initial fit. From here, precision weights are computed without parametric assumptions for use downstream. This approach works well with sufficient read counts, but single cell datasets have substantially few reads per sample, even when using pseudobulk. Modeling the variation in measurement precision due to a Poisson counting process described above and replacing the initial unweighted model fit with precision weights from this Poisson process improves statistical performance specially in the case of low read counts. Because the weights are simply the inverse of the observed counts, there is no additional computational cost.

Current versions of dreamlet $\geq v1.1.23$ use this approach for initial weights for the precision weighted regression models.

**Implicit assumptions of other weighting approaches**
In the previous version of `dreamlet`, samples were weighted by the number of cells observed, denoted here by $n_i$. Following the Poisson model, we show here that this makes unreasonable assumptions about the data.

The vector of precision weights across $K$ samples for one gene is

$$\left(n_1\bar{p}_1\,\bar{l}_1,\ n_2\bar{p}_2\,\bar{l}_2,\ \cdots,\ n_K\bar{p}_K\,\bar{l}_K\right). \quad (17)$$

If we assume that the expression fraction of the given gene is the same in all samples, then $\bar{p}_i$ has the same value for all $i$ denoted here as $p$. Since only the *relative* weights are important, $p$ can be factored out to give the weight vector

$$p\left(n_1\bar{l}_1, n_2\bar{l}_2, \cdots, n_K\bar{l}_K\right) \propto \left(n_1\bar{l}_1, n_2\bar{l}_2, \cdots, n_K\bar{l}_K\right). \tag{18}$$

When the mean library size for each sample are equal, this corresponds to weighting each sample by the number of cells:

$$(n_1, n_2, \cdots, n_K). \tag{19}$$

Therefore, weighting samples by the number of cells makes a number of critical assumptions about the data and generative model of read counts:

1. the expression fraction $\bar{p}_i$ is constant *across* all samples, so there is no expression heterogeneity between samples
2. the mean library size for each sample are equal
3. there is no over-dispersion in counts either within or across samples since a Poisson model is assumed

In practice, these assumptions are not reasonable since the goal of the analysis is to study variation in gene expression across samples.

##### 2.2.2 A negative binomial generative model of read counts

Here, we consider more realistic generative model of read count data. It ends up being impractical for real datasets, but it adds intuition about variation in pseudobulk data.

Instead of counts being drawn from a Poisson distribution with equal rate for all samples, consider a more complex model with variation in Poisson rates. Let the count for a given gene from cell $j$ and sample $i$ be drawn from a Poisson distribution with rate $p_{i,j}l_{i,j}$ where $l_{i,j}$ is the library size for the given cell and $p_{i,j} \in (0,1)$ is the expression fraction from cell $j$ corresponding to the given gene. Recall that over-dispersed count data can be modeled with a negative binomial distribution, which corresponds to a Poisson model of counts with rates drawn from a gamma distribution. Based on this, let the Poisson rate be drawn from a gamma with shape $\alpha_i$ and rate $\beta_i$ that are constant for all cells from sample $i$. The counts are then distributed according to

$$c_{i,j} \sim Pois(p_{i,j}l_{i,j}) \tag{20}$$
$$p_{i,j} \sim \Gamma(\alpha_i, \beta_i). \tag{21}$$

where the $p_{i,j}$ has mean $\mu_i = \alpha_i/\beta_i$ and variance $\sigma_i^2 = \alpha_i/\beta_i^2$, following properties of the gamma distribution. Since the library size is fixed, the distribution of the Poisson rate is

$$p_{i,j}l_{i,j} \sim \Gamma(\alpha_i, \beta_i/l_{i,j}), \tag{22}$$

following properties of scaling gamma random variables. Therefore, $c_{i,j}$ has a negative binomial distribution that is natural for modeling over-dispersed counts.

The expected count is

$$\begin{aligned} E\left[c_{i,j}\right] &= E\left[p_{i,j}l_{i,j}\right] && (23)\\ &= E\left[p_{i,j}\right]l_{i,j} && (24)\\ &= \mu_i l_{i,j} && (25) \end{aligned}$$

and using the law of total variance gives the variance as

$$\begin{aligned} \text{var}\left[c_{i,j}\right] &= E\left[\text{var}\left(c_{i,j}|p_{i,j}\right)\right] + \text{var}\left(E\left[c_{i,j}|p_{i,j}\right)\right]\right) && (26)\\ &= E\left[p_{i,j}l_{i,j}\right] + \text{var}\left(p_{i,j}l_{i,j}\right) && (27)\\ &= \mu_i l_{i,j} + \sigma_i^2 l_{i,j}^2 && (28) \end{aligned}$$

Following the workflow from the previous section, we derive the mean and variances of pseudobulk counts and log CPM. The mean of the pseudobulk count $\tilde{c}_i = \sum_{j=1}^{n_i} c_{i,j}$ is

$$\begin{aligned} E\left[\tilde{c}_i\right] &= E\left[\sum_{j=1}^{n_i} c_{i,j}\right] && (29)\\ &= \sum_{j=1}^{n_i} E[c_{i,j}] && (30)\\ &= \mu_i \sum_{j=1}^{n_i} l_{i,j} && (31)\\ &= \mu_i n_i \bar{l}_i && (32) \end{aligned}$$

The variance is

$$\begin{aligned}
\text{var}\left[\tilde{c}_i\right] &= \text{var}\left[\sum_{j=1}^{n_i} c_{i,j}\right] && (33)\\
&= \sum_{j=1}^{n_i} \text{var}\left[c_{i,j}\right] && (34)\\
&= \sum_{j=1}^{n_i} \left[\mu_i l_{i,j} + \sigma_i^2 l_{i,j}^2\right] && (35)\\
&= \mu_i n_i \bar{l}_i + \sigma_i^2 \sum_{j=1}^{n_i} l_{i,j}^2 && (36)\\
&= \mu_i n_i \bar{l}_i + \sigma_i^2 n_i \bar{\zeta}_i && (37)\\
&= n_i \left(\mu_i \bar{l}_i + \sigma_i^2 \bar{\zeta}_i\right) && (38)
\end{aligned}$$

where $\bar{\zeta}_i = \frac{1}{n}\sum_{j=1}^{n_i} l_{i,j}^2$ is the mean squared library size for sample $i$.

Letting the counts per million (CPM) for pseudobulk computed from sample $i$ be $a_i = 1e6 \cdot \tilde{c}_i/(n_i\bar{l}_i)$, the mean is

$$\begin{aligned}
E\left[a_i\right] &= E\left[1e6 \cdot \tilde{c}_i/(n_i\bar{l}_i)\right] && (39)\\
&= 1e6 \cdot E\left[\tilde{c}_i\right]/(n_i\bar{l}_i) && (40)\\
&= 1e6 \cdot (\mu_i n_i \bar{l}_i)/(n_i\bar{l}_i) && (41)\\
&= 1e6 \cdot \mu_i && (42)\\
\text{var}\left[a_i\right] &= \text{var}\left[1e6 \cdot \tilde{c}_i/(n_i\bar{l}_i)\right] && (43)\\
&= 1e12 \cdot \text{var}\left[\tilde{c}_i\right]/(n_i\bar{l}_i)^2 && (44)\\
&= 1e12 \cdot \frac{n_i\left(\mu_i\bar{l}_i + \sigma_i^2\bar{\zeta}_i\right)}{(n_i\bar{l}_i)^2} && (45)\\
&= 1e12 \cdot \frac{\mu_i + \sigma_i^2\bar{\zeta}_i/\bar{l}_i}{n_i\bar{l}_i} && (46)
\end{aligned}$$

When $\sigma_i^2 = 0$, there is no expression variation with sample $i$ and the variance reduces to

$$\text{var}\left[a_i\right] = 1e12 \cdot \frac{\mu_i}{n_i\bar{l}_i}, \quad (47)$$

which resembles the variance in the Poisson case above.

Now consider the variance of the log CPM approximated using the delta method to give

$$\begin{aligned}
E[\log(a_i)] &\approx \log(E[a_i]) && (48)\\
&= \log(1e6) + \log(\mu_i) && (49)\\
\mathrm{var}[\log(a_i)] &\approx \mathrm{var}(a_i)/E[a_i]^2 && (50)\\
&= \frac{1e12 \cdot \frac{\mu_i + \sigma_i^2 \bar{\zeta}_i/\bar{l}_i)}{n_i \bar{l}_i}}{(1e6 \cdot \mu_i)^2} && (51)\\
&= \frac{\mu_i + \sigma_i^2 \bar{\zeta}_i/\bar{l}_i}{\mu_i^2 n_i \bar{l}_i} && (52)\\
&= \frac{1}{\mu_i n_i \bar{l}_i} + \frac{\sigma_i^2 \bar{\zeta}_i}{\mu_i^2 n_i \bar{l}_i^2}. && (53)
\end{aligned}$$

The variance decomposes into two components. The first represents the variance due to finite read count, number of cells and library size, and matches the variance from the Poisson case above. The second represents the variation in expression among the cells from the same sample. This recapitulates the intuition that increasing expression rate, number of cells and mean library size all reduce variation in log CPM, while increasing within-sample expression variation (i.e. $\sigma_i^2$) increases the variance.

The variance can also be written as

$$\mathrm{var}[\log(a_i)] \approx \frac{1}{\mu_i n_i \bar{l}_i}\left(1 + \frac{\sigma_i^2 \bar{\zeta}_i}{\mu_i \bar{l}_i}\right) \quad (54)$$

where the first factor is the variance when $\sigma_i^2 = 0$, and the second term is the variance inflation due to expression variation among the cells from the same sample. This term does not depend on the number of cells observed.

Yet the lack of a good estimator for the true within-sample expression variance $\sigma_i^2$ currently prevents the use of this generative model in the context of precision-weighted regression models applied here.

##### 2.2.3 Simulations

We performed simulations in order to evaluate the accuracy of these estimators. Simulations were performed with cell count per sample ranging 2 to 1000, library size per cell ranging from 1k to 15k reads, using a gene with $\mu = 0.001$ and $\sigma^2 = 2 \times 10^{-7}$. The empirical estimators based on the delta method approximation of the Poisson generative model accurately estimated variance parameters when there was sufficient read counts per sample (**Supplementary Figure 11A,B**). As the number of cells per sample increases, the variance of the log counts per million decreases (**Supplementary Figure 11C**).

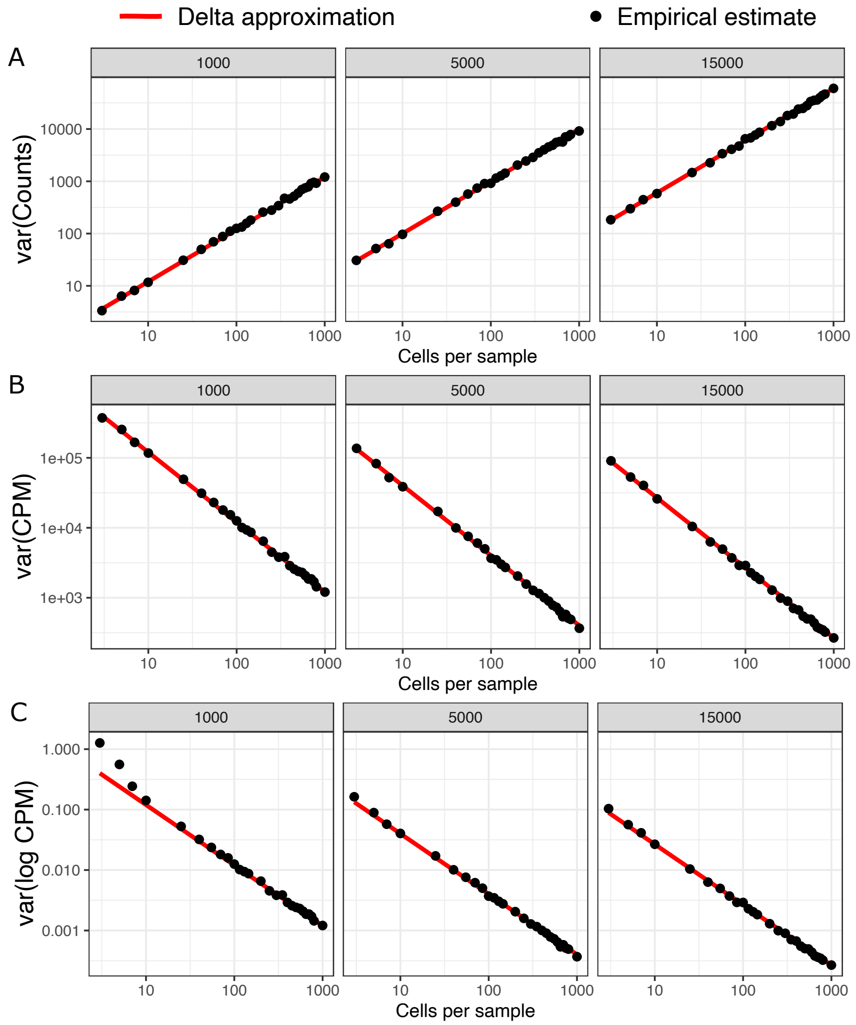

**Supplemental Figure 11: Variance estimates from overdispersed counts**. Delta approximations (red), and empirical estimates from simulated data (black) are shown for multiple variance parameters as a function of the number of cells per sample on the x-axis and the library size per cell on the columns. The true values of $\sigma^2$ were used for the three methods. Values are shown for **A)** variance of the counts, **B)** variance of the counts per million, and **C)** variance of log counts per million.

#### 2.3 Empirical Bayes shrinkage for linear mixed models

For small sample sizes, parameter estimates can have high sampling variance. In seminal work, Smyth (Smyth, 2004) developed an empirical Bayes approach that borrows information across genes to estimate the residual variance. The widely used limma package (Ritchie et al., 2015) fits a linear model for each gene, performs the empirical Bayes step, and then computes a moderated t-statistic with a modified null distribution. In the case of a linear model, Smyth's empirical Bayes method uses a conjugate prior on the residual variances and assumes they are drawn from an inverse gamma distribution (i.e. precisions are drawn from a scaled chi-squared distribution) with parameters estimated from the data. A key value in this calculation is the residual degrees of freedom. In the case of a linear model with $n$ samples and $p$ covariates (including the intercept), the residual degrees of freedom ($df_r$) is $n-p$. However, the case of a linear mixed model used here is more complicated. In this case, we show that the residual variance estimates follow a distribution given by a weighted mixture of $n$ chi-squared random variables, where the weights depend on both the data and the estimated model parameters. We match the expected value of this mixture distribution using a single chi-square and use its degrees of freedom to approximate the $df_r$ of the linear mixed model. Importantly, this method is exact in the case of a linear model, is approximate for linear mixed models with any number of random effects, and the approximation improves with the sample size.

##### 2.3.1 Residual degrees of freedom

In the case of the linear model with

$$y = X\beta + \varepsilon \quad (55)$$

$$\varepsilon \sim \mathcal{N}(0, \sigma^2_\varepsilon) \quad (56)$$

$n$ samples and $p$ predictors (including intercept), the hat matrix $H = X(X^TX)^{-1}X^T$ transforms observed and fitted response values according to $\hat{y} = Hy$ (Hastie and Tibshirani, 1990). Similarly, the residuals are

$$r = y - \hat{y} \quad (57)$$

$$= y - Hy \quad (58)$$

$$= (I - H)y. \quad (59)$$

The residual sum of squares is therefore

$$r^Tr = y^T(I-H)^T(I-H)y. \quad (60)$$

For any vector of normally distributed values $z \sim \mathcal{N}(0,1)$ and a positive semi-definite matrix $A$, the quadratic form is distributed according to a mixture of chi-squared distributions weighted according to the eigen-values of $A$. Formally, $z^TAz \sim \sum_i \lambda_i \chi^2_1$, where $\lambda_i$ values are the eigen-values of $A$.

Since the expected value of the sum of weighted $\chi^2$ random variables is the sum of the weights, then $E[z^T Az] = \sum_i \lambda_i = \mathrm{tr}(A)$.

For the linear model, $A = (I - H)^T(I - H)$ and has $n - p$ eigen-values with value 1, and the rest are 0. Therefore, $\mathrm{tr}(A) = p$.

In general,

$$r^T r/\sigma_e^2 \sim \sum_i \lambda_i \chi_1^2 \tag{61}$$

where the weights $\lambda_i$ are defined by $H$. The unbiased estimate of the residual variance is thus $E[\sigma_e^2] = r^T r/(n - \mathrm{tr}(A))$.

In the case of a linear model, this reduces to the standard theory from linear modeling

$$r^T r/\sigma_e^2 \sim \chi_{n-p}^2. \tag{62}$$

For the linear mixed model, we can use a single $\chi^2$ distribution to approximate the weighted mixture of $\chi^2$. Setting the degrees of freedom to $\mathrm{tr}(A)$ matches the mean of the mixture distribution, while being exact in the case of the linear model. The value of $\mathrm{tr}(A)$ is computed based on

$$\mathrm{tr}(A) = \mathrm{tr}\left[(I - H)^T(I - H)\right] \tag{63}$$

$$= n - 2\mathrm{tr}(H) + \mathrm{tr}(HH^T). \tag{64}$$

##### 2.3.2 Mixture of chi-squares

Let $x$ be a weighted mixture of $k$ $\chi_1^2$ values where $\lambda_i \in [0, 1]$ is the weight of the $i^{th}$ component so that $x \sim \sum_{i=1}^k \lambda_i \chi_1^2$. This distribution has $k$ parameters and an expectation $E[x] = \sum_{i=1}^k \lambda_i$. When all $\lambda_i$ are 0 or 1, $x$ is $\chi_\nu^2$ distributed with $\nu = \sum_{i=1}^k \lambda_i$. In this case, matching the mean of the mixture with a single chi-squared distribution is exact. For arbitrary non-negative values of $\lambda_i$, the distribution of $x$ does not reduce to this simple form, but setting $\nu = \sum_{i=1}^k \lambda_i$ approximates the mixture distribution by matching its mean. We examine the chi-square approximation of the weighted sum of chi-square random variables for increasing values of $k$ (**Supplementary Figure 12A**).

While Smyth uses the $df_r$ for both the EB step and the null of the moderated t-statistic, here we used the approximate $df_r$ just for EB. We use the Satterthwaite or KR method for the null distribution. The $df_r$ of a linear mixed model fit by `lme4::lmer()` can be computed by `variancePartition::rdf.merMod()`.

##### 2.3.3 Simulations

The EB step shrinks the MLE estimates toward the global mean to reduce the impact of genes with extremely high or low residual variance inflating the false positive rate (**Supplementary Figure 12B**). Simulations of $p = 20k$ genes, a discrete covariate with two categories modeled as a random effect, and varying sample sizes indicates that the EB post processing of results from a linear mixed models increases statistical performance (**Supplementary Figure 12C,D**).

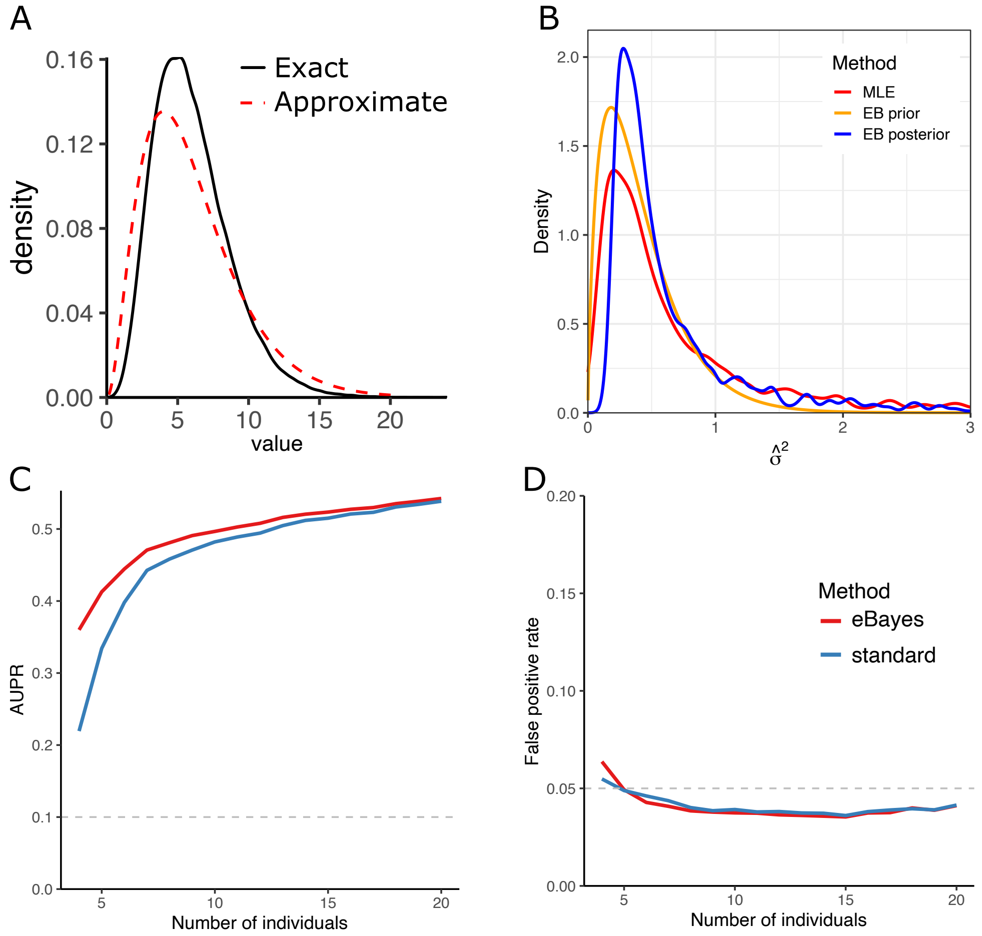

**Supplemental Figure 12: Empirical Bayes shrinkage**. **A)** Approximation of *scaled* chisq distribution with a chisq distribution. Here, 100,000 values were sampled from a weighted mixture of $k$ $\chi^2_1$ variables with $\lambda_i = 0.5$ and the kernel density is plotted in black for 10. The $\chi^2_\nu$ approximation matching the mean is shown in red. **B)** Empirical Bayes (EB) shrinkage of residual variance from simulated dataset of 20k genes with residual precision drawn from $\Gamma(4, 4)$. The EB posterior estimates (blue) shrink the MLE estimates (red) toward the global mean. **C)** Area under the precision-recall curve for simulated data for the standard method (blue) and EB shrinkage (red) for increasing sample size. **D)** False positive rate for these simulations.
